# The C-terminal intrinsically disordered region of a fungal LPMO binds copper and displays anti-fungal properties

**DOI:** 10.1101/2025.11.04.686606

**Authors:** Ketty C. Tamburrini, Koar Chorozian, Clarisse Roblin, Aurore Labourel, Mireille Haon, Sacha Grisel, Evangelos Topakas, Mickael Lafond, Bruno Guigliarelli, Gaston Courtade, Sonia Longhi, Jean-Guy Berrin

## Abstract

Lytic Polysaccharide Monooxygenases (LPMOs) are enzymes that play a crucial role in the degradation of complex polysaccharides such as cellulose and chitin. While LPMOs have attracted significant interest for industrial applications to convert biomass into biofuels, emerging evidence suggests alternative functions in fungal plant pathogenesis, microbial diseases and (micro)organism development. The AA14 LPMO family is widely distributed in filamentous fungi but remains enigmatic as its initially-suspected substrate specificity was recently challenged. In this study, we investigated the disordered C-terminal regions (dCTRs), found in more than half of AA14 family members and hypothesized to have functional relevance. Focusing on the *Pycnoporus coccineus* AA14A LPMO, we used small angle X-ray scattering (SAXS) and circular dichroism to show that its dCTR is highly disordered. It consists of a heavily glycosylated low complexity region followed by a charged C-terminal tail. We uncovered that the *Pco*AA14A dCTR binds copper ions through histidine residues located in the C-terminal tail. Using electron paramagnetic resonance (EPR), we further demonstrated that dimer formation occurs through an oxidative process involving copper and a redox-sensitive cysteine. Noting similarities between the last residues of the C-terminal tail and antimicrobial peptides, we investigated its potential antimicrobial function. We found that the positively charged region of the C-terminal tail selectively inhibits the growth of basidiomycete fungal spores. These data reveal that the dCTRs appended to AA14 LPMOs are functional regions that must be considered to unveil the role of these atypical LPMOs.

**Significance Statement:** Lytic polysaccharide monooxygenases (LPMOs) degrade complex polysaccharides and are central to biomass conversion. Their biological relevance is expanding, with emerging roles in fungal development and host interactions. Despite this broad potential, research has largely focused on their catalytic domain. One area that remains poorly understood is the function of their C-terminal region, predicted to be intrinsically disordered (dCTR). The AA14 LPMO family, whose substrate specificity remains enigmatic, is enriched in dCTRs. This study provides the first experimental evidence of the disordered nature of the dCTR of an AA14 LPMO. Using multidisciplinary approaches, we demonstrate that the dCTR binds metals and exhibits antifungal activity. These findings establish dCTRs as functional regions, essential for understanding the biological roles of LPMOs.

## Introduction

Lytic Polysaccharide Monooxygenases (LPMOs) are enzymes that play a crucial role in breaking down complex polysaccharides, like cellulose and chitin, respectively found in plant and fungal cell walls. Unlike traditional carbohydrate-active enzymes (CAZymes), LPMOs introduce oxidative modifications to these polysaccharides, facilitating their enzymatic degradation by hydrolases (1, 2). LPMOs have an exposed copper atom at their active site, known as Histidine brace (3). Catalysis involves the reduction of the copper ion from Cu(II) to Cu(I) using various electron sources (4). While LPMOs were initially thought to require molecular oxygen, H_2_O_2_ is the most efficient co-substrate for these enzymes (5). Currently, LPMOs are classified into eight families within the auxiliary activities (AA) class of the CAZy database: AA9-AA11 and AA13-AA17 (6). LPMOs are taxonomically widespread, their presence has been confirmed in fungi, bacteria, oomycetes, animals, and even viruses and plants (ferns), expanding our understanding of the ecological significance of these enzymes (7). The substrate specificity of LPMOs, which varies among these AA families, is mostly towards cellulose, chitin, starch or pectin. Despite their widespread abundance, only a few members of the most recently discovered LPMO families have been hitherto characterized and their functions remain unclear. LPMOs have attracted growing interest for various industrial applications, including the production of biofuels, textiles, and nanocelluloses (8, 9). In nature, increasing importance of LPMOs has been emerging in fungal plant pathogenesis (10, 11), diseases caused by bacteria (e.g. pneumonia) (12), fungi (e.g. human meningitis) (13), oomycetes (14) and viruses (e.g. entomopox) (15), as well as in (micro)organism development (16). The AA14 LPMO family was established in 2018, following the biochemical characterization of two LPMOs from the white rot fungus *Pycnoporus coccineus* (syn. *Trametes coccinea*) (17). During growth on plant biomass, AA14 LPMO-encoding genes were found to be up-regulated during fungal growth on wood, and investigations with recombinantly produced *Pco*AA14A and *Pco*AA14B indicated their potential enhancement of enzymatic wood fiber saccharification and activity on heteroxylan covering cellulose fibrils in wood (17). Subsequent research demonstrated synergistic effects when *Pco*AA14B was combined with GH30 xylanases for the degradation of pretreated wood materials (18). However, the substrate specificity of AA14 LPMOs has been recently revisited while studying a new member of the AA14 family, *Tr*AA14A from *Trichoderma reesei* (19). The comparison of this new enzyme with *Pco*AA14A showed that, while these enzymes share many properties with LPMOs, such as the ability to generate and consume H_2_O_2_, they do not seem to act on cellulose-associated xylan, opening several questions on the role of this enigmatic family widespread in filamentous fungi. A recent study hypothesized that AA14 could be involved in the degradation of glucuronoxylomannan (20), which is one of the components of fungal cell wall (21, 22).

Interestingly, the AA14 family is the only LPMO family that does not carry any carbohydrate binding modules (CBMs). Instead, AA14 LPMOs display intrinsically disordered C-terminal regions (dCTRs) (23). Intrinsically disordered proteins (IDPs) or regions (IDRs) defy the classical structure-function paradigm. Contrary to the conventional notion of proteins folding into single, stable three-dimensional structures, IDPs/IDRs are characterized by the lack of a stable secondary and/or tertiary structure (24). Instead, they adopt an ensemble of conformations that dynamically interconvert into each other. The conformational heterogeneity typical of IDPs/IDRs arises from a depletion of hydrophobic residues and an enrichment of charged amino acids (25). IDRs can play key roles in critical cellular processes, including signal transduction, transcriptional regulation, and molecular recognition (26). Their inherent flexibility allows them to interact with multiple binding partners and adapt to different environments, mediating complex and context-dependent interactions within cellular networks (27).

Among all LPMO families, the AA14 family is the most enriched in dCTRs, as they occur in 57% of the sequences. As these regions are usually removed prior to LPMO characterization, their potential contribution to LPMO function remains unknown. The only previously reported functional role for a dCTR comes from an AA9 LPMO from a fungal phytopathogen, where the dCTR was shown to mediate LPMO dimerization both *in vitro* and *in vivo via* a disulfide bridge (11). This dimerization enhances both binding and activity on cellulose. Furthermore, deletion of the gene encoding this AA9 LPMO impaired the formation of the appressorium, a specialized infection structure, and delayed host penetration.

In this study, we focused on the long dCTR appended to an AA14 from *Pycnoporus coccineus*. We combined biochemical and biophysical approaches to experimentally characterize the conformational properties of the dCTR, along with microbiological and microscopy analyses to evaluate its functional role. Our results establish that dCTRs are functional regions that must be considered and further investigated to elucidate the role of these atypical and widely distributed LPMOs.

## Results

### Disordered C-terminal regions (dCTRs) are widely distributed across the AA14 family

The AA14 family is the most dCTR-enriched LPMO family. To explore the dCTR distribution within the family, we conducted an in-depth bioinformatic analysis on 570 AA14 LPMOs. The phylogenetic analysis revealed that dCTRs are present in all clades (**Fig. 1A**) and that wood decaying and mycorrhizal fungi possess a higher number of dCTRs compared to other lifestyles (**Fig. S1**). Although the dCTRs in the various clades exhibit differences in their length, with median values ranging from 41 residues (clade 5) to 120 residues (clade 6) (**Fig. S2A**), the proportion of disordered residues is considerably high as most dCTRs contain over 70% of residues predicted to be disordered (**Fig. S2B**). As observed for the dCTRs from other LPMO families (23), Ser, Thr, Pro, and Ala constitute on average half of the AA14_dCTRs composition. On the other hand, the last 30 residues of AA14_dCTRs are enriched in Arg, His, Phe, Lys, Met, and Trp (**Fig. S2C**). Basidiomycetes possess a higher percentage of LPMOs bearing dCTRs (55% of sequences) in comparison to ascomycetes (30% of sequences) and have a longer median of dCTRs (110 residues) compared to ascomycetes (68 residues) (**Fig. S3A**). However, no significant difference was observed in the fraction of disordered residues (**Fig. S3B**). The clade 8 attracted our interest as (i) it is mainly composed of white rot fungi, (ii) it is particularly enriched in dCTRs, and (iii) most sequences have a highly positively charged patch at the end of the dCTR (**Fig. 1** and **Fig. S2**). Interestingly, the AA14A from the white rot fungus *Pycnoporus coccineus* (*Pco*AA14A) belongs to this clade. While we have previously investigated the properties of *Pco*AA14A catalytic domain (17, 19), its dCTR has been neglected and the protein was recombinantly expressed without its dCTR. The length of *Pco*AA14A_dCTR (137 residues) is close to the median value (115 residues) of dCTRs identified in the AA14 family (**Fig. 1B** and **Fig. S2A**). The 3D structural model of *Pco*AA14A (**Fig. 1C**), as determined using AlphaFold3 (28), features a catalytic domain which is characteristic of LPMOs. Analysis of the pLDDT values (29), revealed low confidence scores for the dCTR (average 43.4%; 80% of atoms <50) suggesting a disordered nature (30). The HCA plot of the dCTR further supports its disordered nature (**Fig. S4**). The alignment of the last residues of the dCTRs from 17 AA14 sequences within the clade of *Pco*AA14A revealed the presence of a cysteine residue conserved in approximately half of the sequences, along with a conserved patch of Arg and His residues at the end of the sequence (**Fig. 1B**). This latter feature, which is reminiscent of the X283 short linear motif (SLiM) identified in AA9 LPMOs (31), could suggest a common function among AA14_dCTRs.

**Figure 1.**
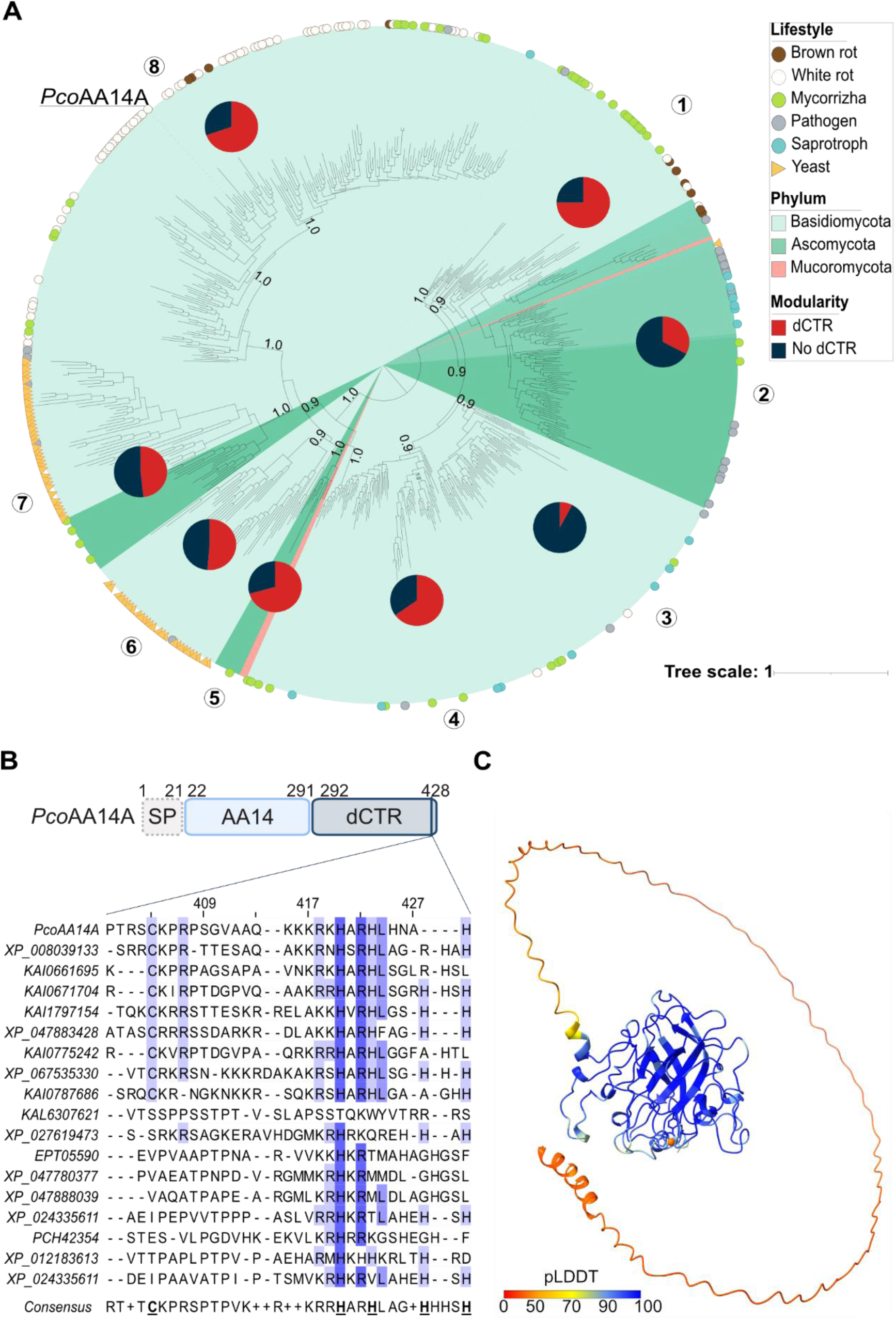
Distribution and nature of disordered C-terminal regions (dCTRs) across the AA14 family. (**A**) Phylogenetic tree of AA14 family. The phylogenetic tree was built based on the catalytic domains of 570 AA14 sequences. The fraction of LPMOs bearing dCTRs within each clade is represented in red in the pie chart. The different clades are numbered. The branching probabilities are indicated when values are equal to or higher than 0.8.) (**B**) Modular organization of *Pco*AA14A and multiple sequence alignment of the last residues within the dCTR along with the consensus sequence from 17 AA14s in the *Pco*AA14A clade, as obtained by aligning the sequences with MCoffee (78). Residues with an identity percentage higher than 20% are colored purple. (**C**) AlphaFold3 model of *Pco*AA14A colored according to pLDDT values. The copper atom coordinated in the active site is shown as a sphere.

### The C-terminal region of *Pco*AA14A is intrinsically disordered, drives enzyme dimerization and enhances protein stability

To experimentally assess the conformational properties of the dCTR of *Pco*AA14A, and in view of getting insights into its functional role, we heterologously produced in the yeast *Pichia pastoris* three *Pco*AA14A forms: the full-length protein (*Pco*AA14A_FL), the catalytic domain alone (*Pco*AA14A_CD) and the dCTR alone (*Pco*AA14A_dCTR). The three proteins were successfully secreted in the *P. pastoris* culture medium and purified to homogeneity. SDS-PAGE analysis of the purified proteins (**Fig. S5A**) revealed that *Pco*AA14A_FL migrates with an apparent molar mass (MM) of 115 kDa, while *Pco*AA14A_CD and *Pco*AA14A_dCTR display an apparent MM of 52 kDa and 70 kDa, respectively. The three apparent MMs are higher than the theoretical ones (46 kDa, 32 kDa and 14 kDa, for *Pco*AA14A_FL, *Pco*AA14A_CD and *Pco*AA14A_dCTR, respectively). Size exclusion chromatography (SEC) coupled to multi-angle laser light scattering (SEC-MALLS) analysis revealed that the three proteins are glycosylated, with glycans representing 34% and 57% of the molar masses in *Pco*AA14A_FL and *Pco*AA14A_dCTR, respectively (**Fig. S6** and **Table S1**). The aberrant electrophoretic migration of *Pco*AA14A_FL and *Pco*AA14A_dCTR likely arises from both glycosylation and the disordered nature and high proline content (10% to be compared to 4.7% for proteins in the UniProtKB/Swiss-Prot database) of the dCTR. Indeed, aberrant electrophoretic migration is a hallmark of IDRs because of their typical compositional bias, e.g., high net charge and low hydrophobicity, and high proline content that causes them to bind less SDS than globular proteins (32). The SEC and SEC-MALLS profiles revealed that while *Pco*AA14A_CD elutes in a single peak, *Pco*AA14A_FL and *Pco*AA14A_dCTR feature an additional smaller peak (i.e. peak 1) beyond a major peak (i.e. peak 2) (**Fig. S5B** and **Fig. S6**). SEC-MALLS analysis revealed that peak 1 is compatible with a dimeric species (**Table S1**), and SDS-PAGE analysis of the dCTR in the absence of DTT enabled detection of an additional faint band with an apparent MM twice that of the species observed under reducing conditions (**Fig. S7**), consistent with a disulfide bridge-driven dimerization. Since no dimerization is observed for the CD alone, this dimerization is probably driven by Cys404 in the dCTR, mirroring our previous findings on an AA9 LPMO (11).

To evaluate the secondary structure content of *Pco*AA14A_dCTR, we used circular dichroism in the far ultraviolet (UV) region (**Fig. 2A**). The *Pco*AA14A_dCTR spectrum displays a large negative peak centered at 198 nm, low intensity in the 220–230 nm region, and low ellipticity at 190 nm, features that are all typical of a disordered protein lacking any stable organized secondary structure (**Fig. 2A**). Indeed, spectral deconvolution revealed a high content (about 80%) of unordered structure and a very low content in secondary structure elements (3% and 8% of α-helices and β-strands, respectively) (**Fig. S8A**). Comparison of the ellipticity values at 200 and 222 nm of *Pco*AA14A_dCTR with those of a set of well characterized IDPs, encompassing both premolten globules (PMGs) and random coils (RCs) (25), revealed that *Pco*AA14A_dCTR falls in the RC region of the plot (**Fig. S8B**). To assess the folding potential of *Pco*AA14A_dCTR, we recorded the far-UV circular dichroism spectra in the presence of increasing concentration of trifluoroethanol (TFE), a secondary structure stabilizer mimicking the hydrophobic environment that proteins experience upon establishing interactions. Therefore, TFE is frequently used to uncover hidden structural propensities in IDPs, i.e. to unveil protein regions that are prone to undergo induced folding (32). The addition of TFE induces only a moderate gain of secondary structure, as judged from (i) the moderate increase in the ellipticity at 190 nm, (ii) the slight reduction in the intensity of the negative peak at 198 nm, and (iii) the lack of appearance of sharp peaks at 208 and 222 nm (**Fig. 2A**). These results indicate that the dCTR is poorly foldable.

**Figure 2.**
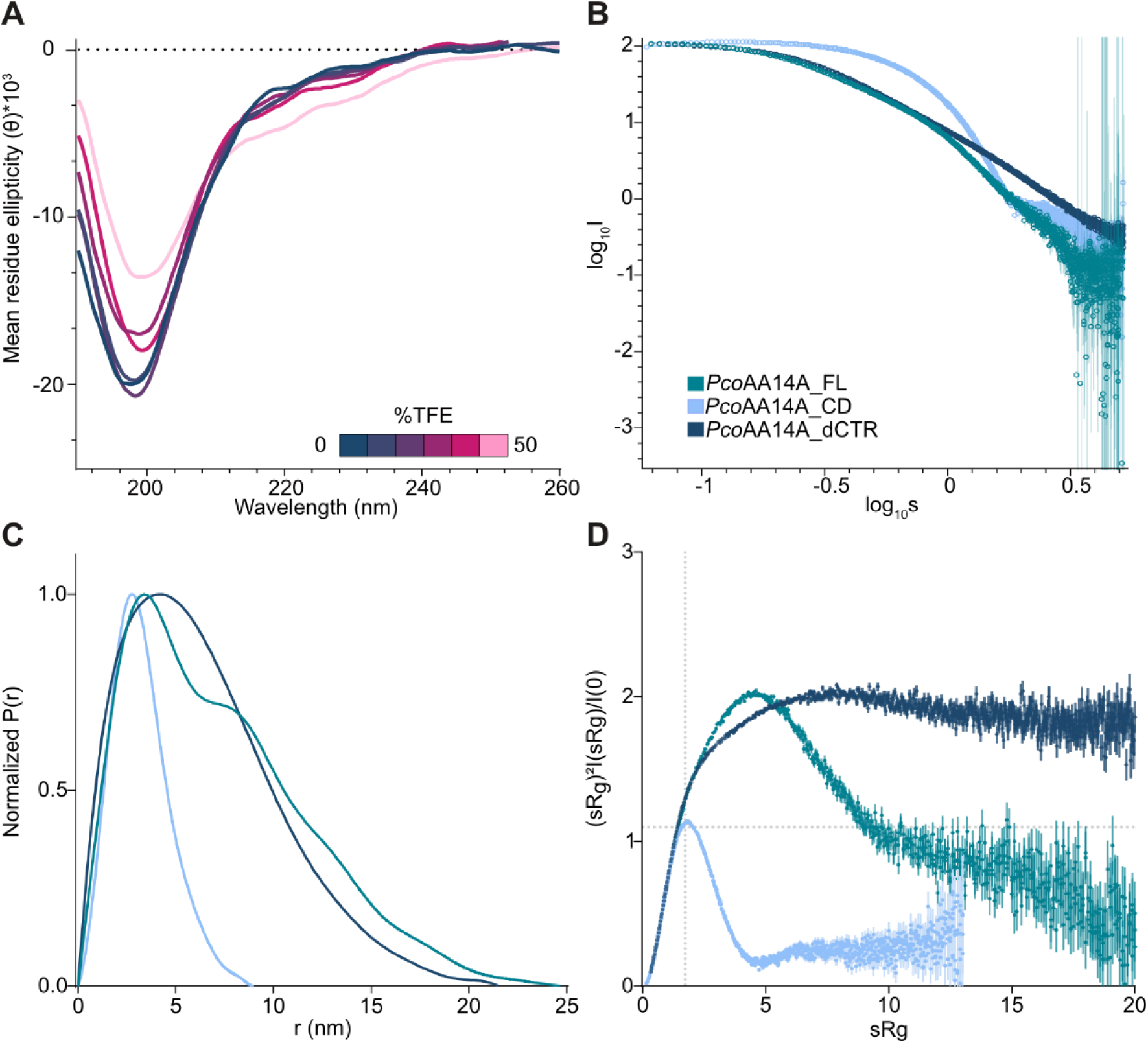
Structural analysis of *Pco*AA14A forms by circular dichroism and SEC-SAXS. (**A**) Far-UV circular dichroism of *Pco*AA14A_dCTR in the presence of increasing concentration of TFE (0-50%). (**B**) Log-Log plots of PcoAA14A_FL (dark green), PcoAA14A_CD (light blue) and PcoAA14A_dCTR (blue). (**C**) Pairwise distance distribution function of *Pco*AA14A_FL (dark green), *Pco*AA14A_CD (light blue) and *Pco*AA14A_dCTR (blue). (**D**) Dimensionless Kratky plot *Pco*AA14A_FL (dark green), *Pco*AA14A_CD (light blue) and *Pco*AA14A_dCTR (blue). The cross-hair lines mark the Guinier-Kratky point (1.73, 1.1), corresponding to the main peak position for globular proteins.

To investigate in more depth the structural properties of *Pco*AA14A, we performed SEC coupled to small-angle X-ray scattering (SEC–SAXS). The three forms show linearity in the Guinier region of the scattering curves (**Fig. S9**), indicating that all samples are monodisperse, thus enabling meaningful estimation of the radius of gyration (R_g_). The derived R_g_ values are 5.78 ± 0.02 nm, 2.50 ± 0.00 nm, and 5.22 ± 0.02 nm for *Pco*AA14A_FL, *Pco*AA14A_CD, and *Pco*AA14A_dCTR, respectively. While the R_g_ value of *Pco*AA14A_CD is close to the one expected for a folded protein of the same length (1.92 nm), as calculated from Flory’s equation (see Mat & Methods), the R_g_ values of *Pco*AA14A_FL and *Pco*AA14A_dCTR are larger than the R_g_ values expected for folded proteins (2.14 Å and 1.56 nm, respectively). The shapes of the pairwise distance distribution functions (P(r)) confirm that both *Pco*AA14A_FL and *Pco*AA14A_dCTR have an elongated conformation, with a maximum intramolecular distance (D_max_) of 24.7 nm and 21.5 nm, respectively, while the bell-shaped P(r) function of *Pco*AA14A_CD suggests a globular conformation with a D_max_ of 9.0 nm (**Fig. 2C**). To assess the flexibility of the three *Pco*AA14A forms, we looked at their dimensionless Kratky plot. The *Pco*AA14A_CD plot displays a typical bell shape with a maximum at around sR_g_ = √3, confirming that the catalytic domain is folded (**Fig. 2D**). The *Pco*AA14A_dCTR plot is characterized by an initial monotonic increase in the low-s region followed by a plateau in the high-s region. This feature, together with a Porod exponent of 2.2, suggests that *Pco*AA14A_dCTR adopts a self-avoiding random walk (SAW) conformation in solution (**Fig. 2D**). Finally, the *Pco*AA14A_FL plot displays an intermediate profile between those of *Pco*AA14A_CD and *Pco*AA14A_dCTR, confirming that the dCTR is highly flexible both in isolation and in the context of the full-length protein. In conclusion, SEC-SAXS data confirm that the dCTR is highly flexible in solution and it is also extended in the full-length protein.

To investigate if the dCTR could affect the thermal stability of the protein, we performed differential scanning fluorimetry (DSF, also called thermal shift assay) and monitored the unfolding curves of *Pco*AA14A_FL and *Pco*AA14A_CD. The two unfolding curves show a single cooperative unfolding step reflecting a two-state transition, typical of proteins with a single folded domain (**Fig. S10**). However, while the *Pco*AA14A_CD melting curve shows a sharp and fast thermal denaturation transition between the folded and unfolded state, typical of globular proteins, *Pco*AA14A_FL shows a slower kinetics, reflecting an unfolding process that takes place over a broader temperature range. The midpoint of the transition curve was calculated using the Boltzmann equation, returning a T_m_^app^ of 45.0 ± 0.5 °C for PcoAA14A_FL and a T_m_^app^ of 41.1 ± 0.1 °C for *Pco*AA14A_CD. The higher T_m_^app^ of *Pco*AA14A_FL suggests that the dCTR stabilizes the protein.

### The disordered C-terminal region of *Pco*AA14A binds copper and undergoes an oxidative-driven dimerization

We next investigated the metal-binding ability of *Pco*AA14A_dCTR. As mentioned earlier, the C-terminal region of *Pco*AA14A_dCTR displays four His, two of which are strictly conserved in a subset of sequences of the clade that also feature a conserved Cys (**Fig. 1B**). The arrangement of the His residues within the AlphaFold3 model of the region encompassing the last 19 residues suggests a potential for metal coordination (**Fig. S11**). To experimentally ascertain whether the dCTR is effectively able to bind metals, we first generated a *Pco*AA14A_dCTR form devoid of the hexa-histidine tag (**Fig. S12A**). Note that circular dichroism studies revealed that the untagged form of *Pco*AA14A_dCTR has the same overall secondary structure content of the tagged one (**Fig. S12B**). A first hint pointing to the ability of the dCTR to bind metals comes from the ability of the untagged form of *Pco*AA14A_dCTR to bind to a HisTrap column, suggesting that it can bind divalent metal ions (**Fig. S12C**). To further investigate this metal binding ability, we carried out inductively coupled plasma mass spectrometry (ICP-MS) analysis. Untagged *Pco*AA14A_dCTR was incubated with copper or zinc and the unbound ions were removed by desalting. ICP-MS data revealed that *Pco*AA14A_dCTR harbors approximately 0.6 copper atom *per* protein molecule and shows only a poor, if any, binding to zinc (0.1 zinc atom per protein molecule). If *Pco*AA14A_dCTR is incubated with both metals, only copper is found (0.4 copper atom *per* protein molecule). Isothermal titration calorimetry (ITC) was then used to obtain further insights into copper binding (**Fig. S13**). The titration of copper into the ITC reaction cell containing *Pco*AA14A_dCTR resulted in a net heat release, confirming that *Pco*AA14A_dCTR binds copper with a stoichiometry of 0.45 copper ions per molecule (**Fig. S13A**), a value in reasonable agreement with ICP-MS data. The apparent equilibrium dissociation constant (K_D_) was estimated to be in the micro-molar range (*K*_D_= 31.3 ± 5.0 µM, n=3). Similar results were obtained by titrating copper into the ITC reaction cell containing a synthetic peptide (P1, **Table S2**) corresponding to the last 19 residues of *Pco*AA14A_dCTR (**Fig. S13B**), confirming that this region is responsible for copper binding.

To further investigate the nature of the interactions of Cu^2+^ ions with the dCTR, P1 was incubated with increasing amounts of Cu^2+^ ions and then monitored by electron paramagnetic resonance (EPR) spectroscopy. A typical axial Cu^2+^ EPR signal develops around g_//_ = 2.242 and g_┴_ = 2.052 with A_//_(^63,65^ Cu) = 18.7 mT and exhibits a superhyperfine structure at g_┴_ arising from the coordination with 3 or 4 ^14^N atoms (A(^14^N) = 1.6 mT) (**Fig. S14**). This signal is not dependent on the used buffer (HEPES or MOPS) and reaches its maximum intensity for 0.5 eq Cu^2+^/P1 in agreement with ICP-MS and ITC results. Because we suspected Cys404 in the dCTR to be involved in enzyme dimerization, we also studied by EPR a longer peptide (P3) encompassing the last 25 residues, and hence starting with a Cys corresponding to Cys404 (**Table S2**). In order to investigate the potential redox role of the dCTR Cys residue, EPR experiments were carried out both in aerobiosis and anaerobiosis. For the series of P3 samples prepared in aerobiosis at low Cu^2+^ concentration (< 0.5 eq), the same Cu^2+^ EPR signal develops as for P1 peptide, and spin intensity was found to be proportional to the amount of added Cu^2+^ (**Fig. 3A**). The superhyperfine structure due to ^14^N nuclei at g┴ = 2.05 is however less resolved indicating a slight broadening of hyperfine lines likely due to an increase of conformation distribution (g– and A-strain broadening) in the P3 peptide compared to P1. Above 0.5 Cu^2+^ eq, the signal broadens and additional components appear revealing the binding of Cu^2+^ ions in other non-specific coordination sites. Strikingly, for the series of P3 samples prepared in anaerobiosis, no Cu^2+^ signal was detected for less than 0.5 eq Cu^2+^ (**Fig. 3B**). This clearly indicates that a significant proportion of Cu(II) ions is reduced to Cu(I) by the P3 peptide in these conditions. The Cu^2+^ signal is observed from 0.5 eq Cu^2+^ and the EPR signal increases linearly with the amount of added Cu^2+^ (**Fig. 3C**). The intercept of the regression line with the horizontal axis gives a value of about 0.35 eq Cu^2+^ showing that about 3 peptides are involved in the reduction of one Cu(II) ion. This is in good agreement with the mechanism proposed previously for the copper catalyzed oxidation of Cys (33) in which one thiolate provides the electron for the reduction of the Cu(II) ion into Cu(I) and the two others are involved in the Cu(I) coordination according to the scheme:

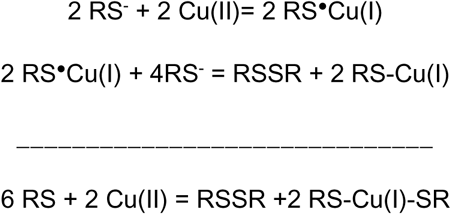

**Figure 3.**
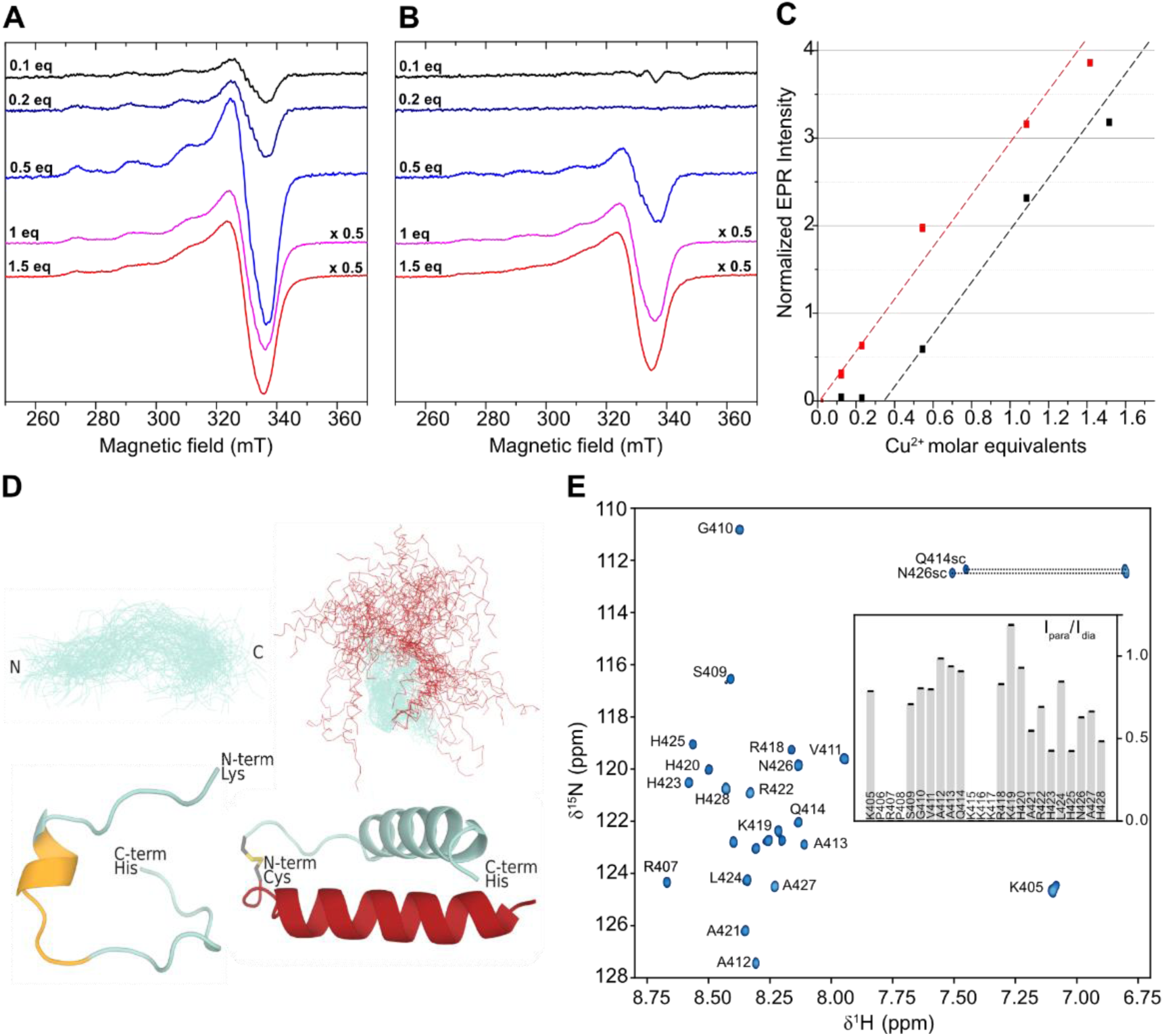
Structural analysis of dCTR model peptides and their binding to copper. EPR analysis of Cu²⁺ interaction with PcoAA14A_dCTR model P3 peptide in aerobiosis (**A**) and anaerobiosis (**B**). EPR conditions: temperature, 60 K; microwave power, 4mW at 9.468 GHz; modulation amplitude, 1 mT at 100 kHz. (**C**) Variation of the Cu²⁺ EPR signal intensity as a function of Cu²⁺ molar equivalents added to P3 peptide in aerobiosis (red squares) or in anaerobiosis (black squares). (**D**) Conformational ensembles of P2 (left) and P3 (right) peptides as obtained from MD simulations and refinement against SAXS data (top) and representative conformations (bottom). The yellow region on the representative conformation of P2 indicates the residues with helical propensity, as obtained using the CSI 3.0 web server (77). (**E**) 2D ^1^H, ^15^N HSQC spectrum of P3 peptide. The sequence-specific residue number assignment of the HN–N crosspeaks is reported next to each peak. Q414sc and N426sc stand for side chains. Residues are numbered based on P3 peptide sequence. Inset: PRE intensity ratio as a function of residue number.

Thus, these results support an oxidative dimerization process of the P3 peptide involving the formation of a disulfide bridge between the unique Cys residues and catalyzed by Cu(II).

To shed light onto the conformational impact of dimerization, we performed both SAXS and molecular dynamics (MD) simulations studies on P3 peptide. As a control, we included in these studies a peptide (P2) encompassing the last 24 residues and thus devoid of Cys (**Table S2**). Conformational ensembles of the P2 and P3 peptides were first generated using 1 µs-long atomistic MD simulations (**Fig S15)** and then refined using SAXS data. The MD trajectories were used to back-calculate ensemble-averaged SAXS curves (**Fig. S16A, B**), where the contribution (weight) of each frame to the scattering profile was optimized iteratively using Bayesian-maximum entropy (BME) approach (34). After BME reweighting the agreement between simulations and SAXS data was good, with reduced χ^2^ values of 1.58 and 1.00 for the P2 and P3 peptides, respectively (**Fig. S15A, B**). The SAXS-refined resulting conformational ensembles, along with representative conformations are shown in **Fig. 3D**. The obtained ensembles illustrate that the P2 peptide is predominantly disordered with a short helical region, whereas P3 is best represented by a disulfide-bridged dimer adopting a mainly helical conformation.

To obtain site-resolved information, the interaction of P3 with copper was also studied by NMR. Backbone resonance assignment of P3 peptide was achieved for most residues, with the N-terminal residue, which usually escapes assignment due to its exchangeable amine group, and the central stretch of three consecutive Lys remaining unassigned (**Fig. 3E**). The narrow chemical shift dispersion in the ^1^H dimension of the ^1^H-^15^N HSQC spectrum – where all signals fall within 7.9-8.7 ppm (**Fig. 3E**) – is consistent with a flexible, disordered conformation. Analysis of chemical shift displacements of the assigned resonances and comparison to chemical shifts of a fully disordered protein, unveiled some helical propensity in the middle of the sequence (**Fig. S17A**), in agreement with circular dichroism data showing that P2 adopts an α-helical structure in the presence of TFE (**Fig. S17B**). Paramagnetic relaxation enhancement (PRE) experiments were performed by titrating Cu²⁺ into the NMR sample. Upon addition of 1.2 molar equivalents of Cu²⁺, significant reduction in signal intensity was observed in the C-terminal region, particularly around the four His residues (**Fig. 3E**, inset). This pattern indicates proximity to the paramagnetic center and supports the presence of a Cu²⁺ binding site in this region. These findings reinforce and extend EPR data that consistently support copper-induced oxidative dimerization of P3 peptide.

### The C-terminal tail of *Pco*AA14A exhibits antifungal properties

To investigate the functional role of the dCTR in *Pco*AA14A, we focused our attention on the positively charged patch found at the end of the dCTR, which is composed of 4 Arg, 5 Lys and 4 His (**Fig. 1B**). Interestingly, the cationic charge of the last 24 residues of *Pco*AA14A_dCTR is reminiscent of some antimicrobial peptides. In particular, we found some sequence similarities with the histatin 5 (Hist-5) peptide, i.e. a human basic salivary peptide with strong fungicidal properties (Helmerhorst et al., 2001). While *Pco*AA14A_dCTR and Hist-5 share only 18% sequence similarity, they both share a high fraction of Lys and His residues (**Fig. S18**). Furthermore, *Pco*AA14A is predicted to be an apoplastic effector protein (with a probability of 51%) by the EffectorP 3.0 predictor (35). Intrigued by these observations, we decided to investigate the antimicrobial activity of *Pco*AA14A_dCTR. We tested Gram-positive and Gram-negative bacteria, as well as ascomycetes and basidiomycetes (**Table S3**). Interestingly, we found that *Pco*AA14A_dCTR negatively impacts the growth of the basidiomycete fungus *Heterobasidion annosum*. The binding of *Pco*AA14A_dCTR to *H. annosum* spores was further tested by a pull-down assay (**Fig. S19**). Quantitative densitometric analysis of the SDS-PAGE revealed that 43 ± 7% of *Pco*AA14A_dCTR is bound to *H. annosum* spores, while no binding was observed for *Pco*AA14A_CD (**Fig. S19**). These results support the ability of *Pco*AA14A_dCTR to bind to *H. annosum* spores. To demonstrate that this activity is due to the positively charged residues of *Pco*AA14A_dCTR, we tested the antimicrobial activity of P1 and P2 peptides (**Table S3**). Interestingly, these peptides showed the same activity on *H. annosum* as *Pco*AA14A_dCTR (**Table S3**), thus confirming that the observed effect is indeed due to the positively charged stretch. We confirmed this antifungal propensity of P1 and P2 peptides against another basidiomycete fungus inhabiting a similar ecological niche, *Coniophora puteana,* and showed that both peptides exhibit potent activity with minimal inhibitory concentrations (MICs) in the micromolar range (**Table S3**).

To further investigate the interaction of peptides with fungal spores and assess their potential impact on spore integrity, we used fluorescence microscopy together with a 5-FAM–labeled P3 peptide. A clear binding to *H. annosum* was observed at the surface of spores and hyphae **(Fig. 4A**). Interestingly, peptide binding remained limited to spores when new hyphae developed (**Fig. 4A**). Consistently with the MICs obtained with P1 and P2, P3 binding also appeared to be dose-dependent, with increased penetration into spores being observed with increasing P3 concentrations (**Fig. 4B**). Strikingly, copper addition induced extra bud-like structures (**Fig. 4C**). No binding of the peptide to *Aspergillus niger* spores was observed (**Fig. 4D**) in agreement with the lack of antimicrobial effect of the dCTR and derived peptides on ascomycete fungi (**Table S3**). To probe the role played by the charged patch, binding experiments were also carried out with a 5-FAM–labeled mutant peptide (P4) in which the KKKRK amino acid stretch was replaced with a charge-neutral AAAAA stretch (**Table S2**). Lack of P4 binding to spores demonstrates the importance of the positively charged patch (**Fig. 4E**). Moreover, high salt concentration abolished peptide binding (**Fig. 4F**), which suggests that the peptide interacts with fungal spores via ionic interactions. Finally, the nucleic acid staining dye propidium iodide revealed that peptide binding was associated with membrane damage and spore loss in *H. annosum*, with this effect being enhanced by copper (**Figure S20**). These findings indicate that the C-terminal tail of *Pco*AA14A exhibits antifungal properties. Binding to spores seems to be mediated by the positively charged region that selectively inhibits the growth of basidiomycete fungal spores, reflecting a membrane-acting mode of action shared with several classes of antimicrobial peptides (36).

**Figure 4.**
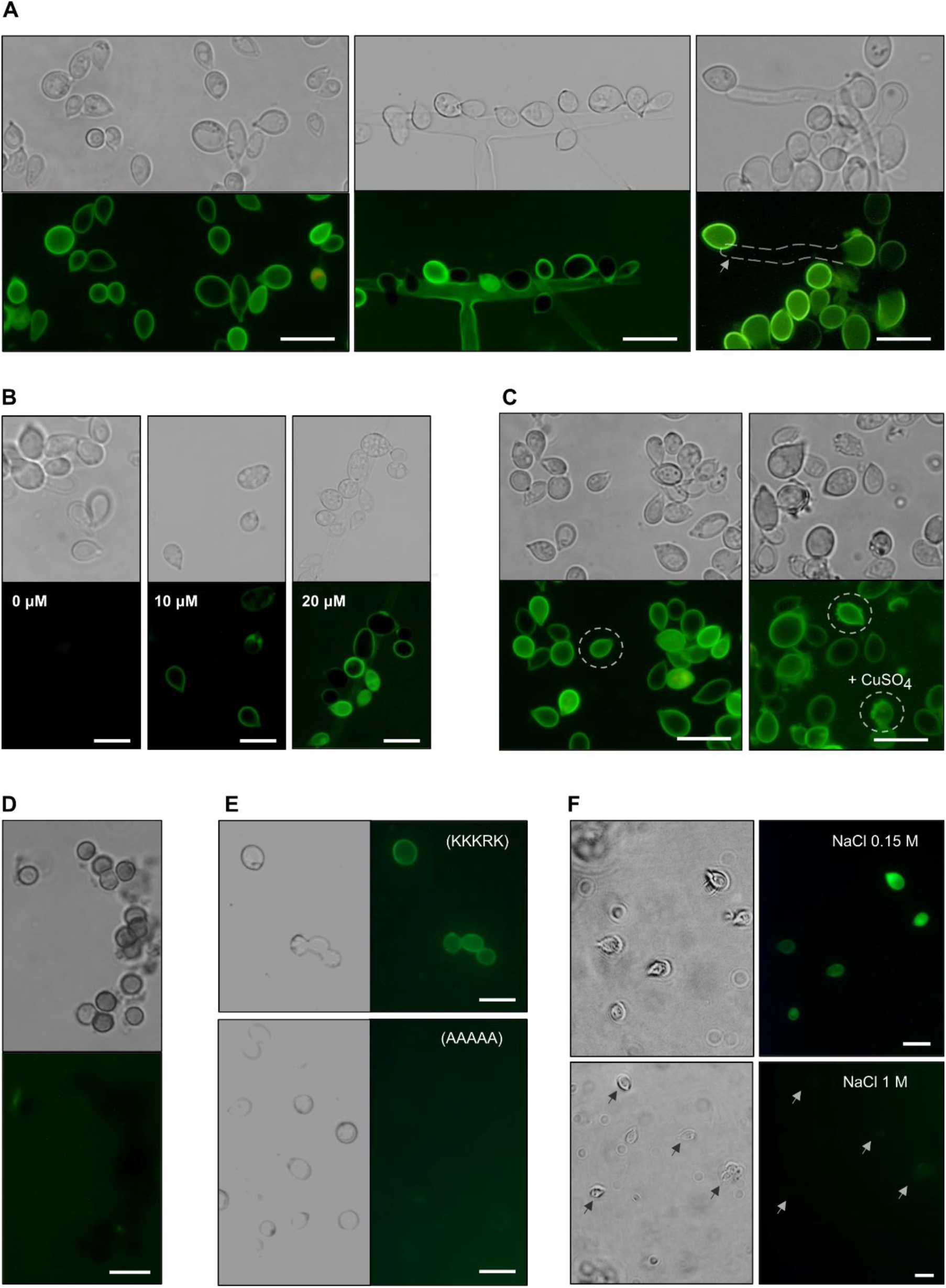
Fluorescence microscopy analysis of dCTR peptides binding to fungal spores. (**A**) *Heterobasidium annosum* incubated with 5-FAM–labeled P3 peptide. Left panel: spores. Central panel: spores and hyphae. Right panel: spores incubated overnight enabling them to develop into hyphae. The brightfield image (top) shows the spores, while the fluorescence image (bottom) reveals the peptide binding to the spore membranes, indicated by the green fluorescence. The absence of green fluorescence in the newly formed hyphae (arrow in the right bottom panel) suggests that P3 remains localized into the spores and does not diffuse into the newly formed hyphae. (**B**) *H. annosum* spores treated with increasing concentrations of 5-FAM-labeled P3 (0, 10 and 20 μM). (**C**) Spores of *H. annosum* incubated with 5-FAM–labeled P3 peptide incubated in the absence (left panel) or presence (right panel) of CuSO_4_. Additional features (more bud bodies) observed in the presence of CuSO_4_ are indicated by dashed lines. (**D**) *Aspergillus niger* spores incubated with 5-FAM–labeled P3 peptide. The brightfield image (top) shows the spores, while the fluorescence image (bottom) indicates a total lack of green fluorescence. (**E**) *H. annosum* spores incubated with 5-FAM–labeled P3 (wild type, KKKRK, top) and P4 (AAAAA, charge-neutral mutant, bottom) peptides. (**F**) *H. annosum* spores incubated with 5-FAM–labeled P3 in HEPES buffer (20 mM, 150 mM NaCl; upper panel) or high-salt concentration buffer (1 M NaCl; bottom panel). Scale bars: 10 μm.

## Discussion

The functional characterization of LPMOs remains challenging. The AA14 family is no exception as its initially-suspected substrate specificity was recently challenged and revisited (19). To solve the puzzle of this enigmatic LPMO family, we decided to focus on the dCTRs of AA14 LPMOs, found in more than half of AA14 members. Our experimental observations revealed high disorder content of *Pco*AA14A_dCTR, both in isolation and in the context of the full-length protein. Circular dichroism studies in the presence of TFE revealed poor foldability of *Pco*AA14A_dCTR, which suggests that the dCTR may function mainly as an entropic chain, where function directly depends on the flexibility and the plasticity of the backbone. Our data suggest that *Pco*AA14A_dCTR consists of two parts with distinct properties: a long low complexity region, enriched in Ser and in predicted O-glycosylation sites, and a functional C-terminal basic region encompassing the last 25 residues. Glycosylation of the low complexity region of the dCTR may further enhance chain expansion and overall flexibility, as observed in other IDRs (37, 38).

The C-terminal region of *Pco*AA14A_dCTR is reminiscent of Hist-5 peptide, an antifungal peptide naturally occurring in the salivary glands of humans and other non-human primates (39). Hist-5 is a polycationic, histidine-rich peptide that affects membrane structure by binding to membrane phospholipids. It has the ability to bind to metal ions (Cu^2+^ and Zn^2+^), a property that has a direct impact on its antimicrobial activity (40–42). Similarly, we have shown that *Pco*AA14A_dCTR binds copper and inhibits the growth of basidiomycetes spores, and that the C-terminal peptide binds and damages *H. annosum* spores. The inherent helical propensities of the C-terminal peptide suggest that it could adopt an α-helical structure in a membrane-mimicking environment, as reported for Hist-5 (43). In addition, its antifungal activity is electrostatically driven and dependent on copper as Hist-5, suggesting some similarities in the antimicrobial mechanism whose precise determinants remain to be elucidated. In a natural context, we hypothesize that *Pco*AA14A_dCTR could be an effector used by *P. coccineus* to compete with other fungi for colonization on woody biomass. The wood-decaying fungi *H. annosum* and *C. puteana* inhabit the same ecological niche of *P. coccineus* and it is well known that direct combat occurs between fungal species during wood decay (44). It has been reported that *P. coccineus* is able to overgrow *H. annosum* and replace *C. puteana*, although the underlying molecular mechanisms have remained elusive so far (45). Our findings provide a likely mechanistic framework to explain this behavior, which could confer a selective advantage to the fungus during wood colonization.

We have also provided the first experimental evidence that the disordered C-terminal tail of an AA14 LPMO is able to bind copper with low affinity. IDRs have been already reported to bind metal ions with lower affinities compared to classical metalloproteins, thus enabling highly dynamic metal-IDR interactions (46). In nature, ectomycorrhiza and pathogenic fungi must tightly regulate copper levels to ensure proper protein functions while avoiding toxicity (47). Ectomycorrhiza fungi play a significant dual role in regulating metal levels by improving mineral nutrition of the host plants and increasing tolerance towards toxic metals. In this context, metal transporters and metalloenzymes are highly expressed in symbiotic hyphae (48). Interestingly, transcriptomic data from *Laccaria bicolor* and *Tuber magnatum* confirm that AA14s are expressed during mycorrhiza establishment and upregulated in fruiting bodies development (49–51). In the same context, the recently-discovered LPMO-like X325 from *Laccaria bicolor* is also upregulated during both ectomycorrhizal and fruiting body formation (52). This protein is structurally similar to LPMOs but lacks the redox activity associated with LPMOs and is anchored to the fungal cell wall through a glycosylphosphatidylinositol (GPI) anchor. In the human pathogen *Cryptococcus neoformans*, a copper binding protein that belongs to the X325 family, was found to regulate copper homeostasis by binding and releasing copper from the cell wall in response to oxidative signaling and metal availability (53). Interestingly, the unique AA9 LPMO from *C. neoformans* (*Cn*Cel1) displays a dCTR (13, 54). Targeted mutation of the *Cn*Cel1 gene revealed that it is required for the expression of stress response phenotypes, including thermotolerance, cell wall integrity, and efficient cell cycle progression (13, 54). However, the actual substrate and activity of *Cn*Cel1 remain to be identified.

*C. neoformans* also possesses a single AA14, which harbors a dCTR. The gene encoding this AA14 exhibits a baseline expression in multiple conditions (e.g. response to changes in pH, as well as in two mutants impaired for growth at high pH during pathogenesis) (55). Altogether, these observations suggest that AA14_dCTRs could play a role in the developmental stages of fungi. The dCTR could mediate the enzyme spatial anchoring and its high flexibility could confer a long reach to the AA14. Moreover, dimerization of AA14 LPMOs, mediated by the dCTR and induced by copper, could enlarge the protein anchoring abilities by mimicking multi-modularity, a property known to provide beneficial effects, including stronger binding to substrates as we recently proposed in the case of AA9 LPMOs (11). Unfortunately, the underlying enzymatic activity of AA14 LPMOs, if any, still remains enigmatic. A recent study on *Sporothrix epigloea* that feeds on the secreted polysaccharide matrix of *Tremella fuciformis*, hypothesized that AA14 could act on fungal cell wall glucuronoxylomannan (20), but no biochemical data are available.

In conclusion, we herein provide evidence for the disordered nature and the unique biochemical properties of an AA14_dCTR and gathered data suggesting an important biological role of its C-terminal tail. This region confers several functions to AA14 LPMOs: (i) high flexibility, (ii) protein stability, (iii) copper binding and copper-induced enzyme dimerization, (iv) ability to bind to fungal spores and (v) antifungal properties. The present results demonstrate that the dCTRs appended to LPMOs are functional and important regions that must be considered to unveil the role of non-canonical LPMOs.

## Materials and Methods

### Bioinformatic analysis

The phylogenetic tree was built as described in Tuveng et al. (2023) (19). Sequences were retrieved by BlastP searches against the protein sequences in GenBank and the JGI MycoCosm databases using *Pco*AA14A and *Pco*AA14B as queries. The CD-HIT Suite (56) was used to remove redundancy by selecting one representative sequence from a cluster of sequences with > 80% sequence identity. Sequences corresponding only to the catalytic domains were aligned using Clustal Omega (57). A phylogenetic tree was built using the PHYML software (58), and the substitution model was selected by Smart Model Selection (SMS) using Akaike Information Criterion (AIC) as statistical criteria. Branch support was calculated using the fast likelihood-based method (aLRT SH-like). The tree was visualized using the Interactive Tree of Life (iTOL) software (59). AA14 sequences with a C-terminal region of unknown function longer than 30 residues were extracted and analyzed using IUPred2A (60). A residue is considered as disordered if the predictor returns a score equal or higher than 0.5. A C-terminal sequence is considered to be disordered (and hence classified as dCTR) if it consists of at least 20 consecutive residues predicted as disordered. The fraction of disordered residues is calculated by dividing the number of residues predicted to be disordered by the length of the dCTR. Only dCTRs having a fraction of disordered residues of at least 50% were considered in the analysis. Amino acid frequencies in each clade of the phylogenetic tree were calculated as the number of a given amino acid in the clade divided by the total number of amino acids in the same clade. For the last 30 amino acids in the dCTRs, amino acid enrichment and depletion in each clade was measured normalizing (i.e. subtracting and then dividing) by the amino acid frequencies of the full-length dCTR. Hierarchical clustering of amino acid enrichment was performed using the gplots package (v3.1.3) (61) and RStudio (RStudio Team (2022). RStudio: Integrated Development Environment for R. RStudio, PBC, Boston, MA URL http://www.rstudio.com/).

### Cloning and production of PcoAA14A FL, CD and dCTR

The proteins were produced using the in-house 3PE platform (*Pichia pastoris* Protein Express; www.platform3pe.com) (62). The nucleotide sequence coding for the *Pco*AA14A_CD, *Pco*AA14A_FL and *Pco*AA14A_dCTR from *Pycnoporus coccineus* (Genbank ID #KY769369) was codon optimized for expression in *Pichia pastoris* (syn. *Komagataella phaffii*) and synthesized (GenScript, Piscataway, USA). Each gene was inserted into the *p*PICZαA expression vector (Invitrogen, Cergy-Pontoise, France). This vector drives the expression of the gene of interest under the control of the AOX1 promoter that is inducible by methanol. The constructs were designed to encode C-terminally His tagged forms. In the case of *Pco*AA14A_dCTR, an additional construct encoding a form devoid of His Tag was also generated. *Pme*I-linearized *p*PICZαA recombinant plasmids were used to transform by electroporation competent *P. pastoris* SuperMan5 cells. Zeocin-resistant transformants were then screened for protein production. The best-producing transformants were grown in 2 L of BMGY medium (10 g.L^-1^ yeast extract, 20 g.L^-1^ peptone, 3.4 g.L^-1^ yeast nitrogen base without amino-acids, 10 g.L^-1^ ammonium sulfate, 10 g.L^-1^ glycerol, 100 mM potassium phosphate buffer pH 6.0, biotin 0.4 mg.L^-1^) in non-baffled flasks at 30°C in an orbital shaker (200 rpm) for 16 h to an OD_600_ of 2 to 6. Expression was induced by transferring cells into 400 mL of BMMY media (10 g.L^-1^ yeast extract, 20 g.L^-1^ peptone, 3.4 g.L^-1^ yeast nitrogen base without amino-acids, 10 g.L^-1^ ammonium sulfate, 100 mM potassium phosphate buffer pH 6.0, biotin 0.4 mg.L^-1^). The medium was supplemented (initially and every day) with 3% (v/v) methanol at 20°C in an orbital shaker (200 rpm) for another 3 days. The cells were harvested by centrifugation (5000 rpm, 10 min at 4°C), and just before purification the pH of the supernatant was adjusted to 7.8 with sodium hydroxide and the supernatant was filtered at 0.45 µm (Millipore, Burlington, Massachusetts, USA).

### Purification of *Pco*AA14A proteins

Filtered and pH-adjusted culture supernatants were loaded onto a 5 mL HisTrap HP column (Cytiva, Illkirch, France) equilibrated with buffer A (Tris-HCl 50 mM pH 7.8, NaCl 150 mM, imidazole 10 mM) that was connected to an Äkta purifier 100 (Cytiva). His-tagged recombinant proteins were eluted with buffer B (Tris-HCl 50 mM pH 7.8, NaCl 150 mM, imidazole 500 mM). Fractions containing recombinant *Pco*AA14A_FL, or *Pco*AA14A_CD or *Pco*AA14A_dCTR were pooled, concentrated using Vivaspin protein concentrators with a 10 kDa cut-off, and dialyzed against 50 mM MES, 150 mM NaCl buffer pH 6.5. Protein concentrations were determined by absorption at 280 nm using a Nanodrop ND-2000 spectrophotometer (Thermo Fisher Scientific, Illkirch, France) and extinction coefficients (*Pco*AA14A_CD: 63,090 M^-1^ cm^-1^, *Pco*AA14A_dCTR: 4,470 M^-1^ cm^-1^; *Pco*AA14A_FL: 67,560 M^-1^ cm^-1^) determined with ProtParam (Expasy). The culture supernatant of *Pco*AA14A_dCTR without His-Tag was loaded onto a HiPrep 26/10 Desalting column (GE Healthcare, Bus, France) equilibrated with buffer A (MES 50 mM pH 6.5, NaCl 10 mM) that was connected to an Äkta pure 100 (Cytiva). Fractions containing the proteins were pooled and loaded onto a 20 mL HiPrep SP FF 16/10 column and recombinant proteins were eluted with buffer B (MES 50 mM pH 6.5, NaCl 1M). Fractions containing recombinant *Pco*AA14A_dCTR were pooled, concentrated as described above, and purified by SEC, using a HiLoad 16/600 Superdex 75 pg column (Cytiva, Illkirch, France) operated at 1 mL min^-1^ and using MES 50 mM, NaCl 150 mM pH 6.5 or sodium acetate 50 mM pH 5.2 as elution buffer. Protein concentrations were determined as described above using an extinction coefficient (*Pco*AA14A_dCTR: 4470 M^-1^.cm^-1^) as determined with ProtParam (Expasy). Protein purity was checked by 4-20% Tris-Glycine precast SDS-PAGE (BioRad, Gemenos, France) analysis, using Coomassie Blue as staining.

### Peptides synthesis

Synthetic peptides, both unlabeled and fluorescently labelled, were all purchased from GenScript Corporation with 95% purity.

### Differential Scanning Fluorimetry (DSF)

Determination of T_m_^app^ was made using a protein thermal shift assay (Thermo Fisher Scientific, Waltham, MA, USA). This assay allows determining T_m_^app^ by monitoring the fluorescence resulting from binding of the SYPRO Orange dye to hydrophobic protein regions, generated by partial or full protein unfolding. SYPRO Orange 5000x was diluted to 5x in buffer MES 50 mM, NaCl 150 mM, pH 6.5 and mixed with *Pco*AA14A_FL or *Pco*AA14A_CD (10 µM). Reactions were heated from 25 to 99 °C in 96-well plates over the course of 60 min in a CFX Connect real-time PCR machine (Bio-Rad, Hercules, CA, USA). The fluorescence was measured using excitation and emission wavelengths of 485 nm and 625 nm, respectively. The experiments were performed in three replicates.

### Circular Dichroism

Far-UV circular dichroism spectra were measured using a Jasco J-810 dichrograph (Hachiōji, Japan), flushed with N_2_ and equipped with a Peltier thermoregulation system set at 20 °C. One-mm thick quartz cuvettes were used. Proteins concentrations were 0.1 mg mL^−1^. Spectra were measured between 260 and 190 nm with a scanning speed of 50 nm/min and a data pitch of 0.2 nm. Response time was set to 4 s and the bandwidth to 2 nm. Spectra were recorded in 10 mM sodium phosphate pH 6.6. Each spectrum is the average of five acquisitions. The spectrum of the buffer was subtracted from the protein spectrum. Spectra were also recorded in the presence of increasing concentrations of TFE (from 10 to 50% v/v). Mean molar ellipticity values per residue (MRE) were calculated as [θ] = 3300mΔA/lcn where l is the path length in cm, n is the number of residues, m is the molecular mass in Daltons and c is the concentration of the protein in mg mL^−1^. Numbers of amino acid residues are 137 for untagged *Pco*AA14_dCTR and 143 for *Pco*AA14A_dCTR. Molecular masses are 13,494 for untagged *Pco*AA14_dCTR and 14,397 Da for *Pco*AA14A_dCTR. The DICHROWEB website (http://dichroweb.cryst.bbk.ac.uk/html/home.shtml) (63), was used to analyze the experimental data in the 190–260 nm range using subtracted spectra. The content in the various types of secondary structure was estimated using the CDSSTR deconvolution algorithm with the reference protein set 7.

### Small-Angle X-ray Scattering (SAXS)

SAXS data were collected at the European Synchrotron Radiation Facility (ESRF, Grenoble, France) and at SOLEIL (Gif-sur-Yvette, France) as described in Table 1.

**Table 1.** SAXS data acquisition parameters.

| Data acquisition |  |  |  |  |  |
| --- | --- | --- | --- | --- | --- |
| European Synchrotron Radiation Facility<br>(Grenoble, France)<br>Beamline BM29 |  |  |  | SOLEIL Synchrotron<br>(Gif-sur-Yvette, France)<br>Beamline Swing |  |
| X-rays wavelength (Å) | 0.992 |  |  | 1.033 |  |
| Energy (keV) | 12.5 |  |  | 12 |  |
| Detector type | Pilatus3 2M in vacuum | Pilatus 1M | Pilatus3 2M in vacuum | Dectris EIGER 4M |  |
| Sample-to-detector distance (m) | 2.867 |  |  | 2.0 |  |
| Temperature (°C) | 20°C |  |  | 20°C |  |
| Gel filtration column | BioSEC 300<br>4.6 x 300 | Superdex 200<br>3.2/300 Increase | BioSEC 300<br>4.6 x 300 | Batch experiments |  |
| Flow rate (mL min<sup>-1</sup>) | 0.25 | 0.075 | 0.25 |  |  |
| s-range | 0.063- 5.209 nm<sup>-1</sup> | 0.038-5.18 nm<sup>-1</sup> | 0.063- 5.209 nm<sup>-1</sup> | 0.13-5.54 nm<sup>-1</sup> | 0.06-5.54 nm<sup>-1</sup> |
| Samples |  |  |  |  |  |
|  | *PcoAA14A_dCTR* | *PcoAA14A_CD* | *PcoAA14A_FL* | P2 | P3 |
| Concentration (mg mL<sup>-1</sup>) | 5.3 | 5 | 2.9 | 2 | 2 |
| Sample volume (µL) | 50 | 50 | 50 | 50 | 50 |
| Buffer | 50 mM MES, 150 mM NaCl pH 6.5 | 50 mM sodium acetate pH 5.2 | 50 mM MES, 150 mM NaCl pH 6.5 | 20 mM Hepes pH 7.2, 100 mM NaCl | 20 mM Hepes pH 7.2, 100 mM NaCl |

Data reductions were performed using the established procedure available at the BM29 beamline or using the software FOXTROT according to the SWING beamline procedure. Buffer background runs were subtracted from sample runs using CHROMIX in the ATSAS program package (Manalastas-Cantos et al., 2021). The final scattering intensities were analyzed using the ATSAS program package. Linearity in the Guinier region was used to exclude sample aggregation. The few data points differing from the Guinier fitting at low angles were deleted and the useful data range was determined with SHANUM (64). The radius of gyration (R_g_) was estimated at low angles (q < 1.3/R_g_) according to the Guinier approximation (65, 66):

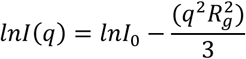

The pairwise distance distribution function, P(r), from which the D_max_ and the R_g_ were estimated, was computed using GNOM and manually adjusted until a good CorMap *p*-value (α > 0.01) was obtained (67). The theoretical R_g_ value (in Å) expected for various conformational states was calculated using Flory’s equation:

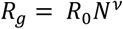

where N is the number of amino acid residues, R_0_ a constant and ν a scaling factor. For IDPs, R_0_ is 2.54 ± 0.01 and ν is 0.522 ± 0.01 (68), for chemically denatured (U) proteins R_0_ is 1.927 ± 0.27 and ν is 0.598 ± 0.028 (68), and for natively folded (NF) proteins R_0_ = √(3/5) × 4.75 and ν = 0.29 (69). The flexibility of the proteins was assessed with the dimensionless Kratky plot ((qR_g_)^2^ I(q)/I_0_ *vs* qR_g_).

### SEC-MALLS

The molecular mass was calculated using analytical size-exclusion chromatography (SEC) on a high-performance liquid chromatography (HPLC) system (Waters) coupled with online multi-angle laser light scattering, ultra-violet light absorbance and refractive index detectors (MALLS/UV/RI) (Wyatt Technology, Santa Barbara, CA, USA). SEC was performed on a Superdex200 10/300 GL Increase column with 50 mM MES pH 6.5, 150 mM NaCl as the elution buffer. The molecular mass was calculated using ASTRA V software (Wyatt Technology) with a refractive index increment (dn/dc) of 0.185 mL/g for protein samples. In the ‘protein conjugate’ analysis the refractive index increment of glycans was set to 0.136 mL/g.

### ICP-MS

Copper content was analyzed using ICP-MS as described in (17). Untagged *Pco*AA14A_dCTR was mineralized, then diluted in ultrapure water, and analyzed by an ICAP Q apparatus (Thermo Electron, Les Ullis, France). The copper concentration was determined using Plasmalab (Thermo Electron) software, at m/z = 63 with an accuracy of ±5%.

### Isothermal titration calorimetry

The binding of *Pco*AA14A_dCTR or P1 to copper was further characterized by isothermal titration calorimetry (ITC) using a MicroCal iTC200 instrument (Malvern). Experiments were carried out at 20°C in a solution containing 20 mM HEPES, 150 mM NaCl, and 2.8 mM glycine (pH 7.4). *Pco*AA14A_dCTR or P1 concentration in the cell was 250 μM whereas the CuSO_4_ concentration in the syringe was 800 μM. Heats of dilution were measured by injecting the ligand into the protein solution. Titration curves were fitted by using MicroCal Origin software, and enthalpy changes (ΔH), dissociation equilibrium constants (K_D_), and stoichiometry were extracted.

### EPR analysis

For the analysis of interactions of *Pco*AA14A_dCTR model peptides with copper, a series of EPR samples was prepared by incubating a solution of either P1 (300 µM) or P3 (400 µM) peptides in 20 mM HEPES (P1) or MOPS (P1, P3) buffer at pH 7.2 with different amounts of CuSO_4_ solution. To minimize the influence of dilution, small volumes of 1 to 5 mM CuSO_4_ solution were directly added to 160 µL samples of peptide solution into EPR tubes, incubated for 5 min and the tubes were frozen in liquid nitrogen for subsequent studies. For the study in anaerobiosis, the same protocol was performed in glove box ([O_2_] < 1.2 ppm) with carefully degassed P3 and CuSO_4_ solutions. EPR measurements were performed on a Bruker ELEXSYS E500 spectrometer equipped with an ER4102ST standard rectangular Bruker EPR cavity fitted to an Oxford Instruments ESR 900 helium flow cryostat. Spin intensity measurements were performed by double integration of EPR spectra recorded in non-saturating conditions.

### Atomistic MD simulations

The a99SB-disp force field was used in combination with GROMACS 2022.2 (https://www.gromacs.org/about.html) (70, 71) to simulate P2 and P3 peptides. The simulation box was filled with water beads, and the system was neutralized with beads corresponding to Na^+^ and Cl^−^ ions. The complex was energy-minimized using a steepest-descent algorithm (50000 steps, 0.01 nm maximum step size) prior to being relaxed in two steps. First, for 100 ps with a time step of 2 fs, using the velocity-rescale thermostat and Verlet cutoff scheme. Second, for 100 ps with a time step of 2 fs, using the Nose-Hover thermostat, Parrinello-Rahman barostat and Verlet cutoff scheme. Simulations were performed on the relaxed models with a time step of 2 fs, using the Nose-Hoover thermostat, Parrinello–Rahman barostat, and Verlet cutoff scheme in the isothermal– isobaric (NPT) ensemble. Data and scripts used for running and analyzing the simulations are available from https://github.com/gcourtade/papers/tree/master/2025/AA14-peptides.

### Calculating SAXS profiles from conformational ensembles

Previous to the analysis, SAXS data for the P2 and P3 peptides at 2 mg mL^−1^ were logarithmically rebinned with a bin factor of 1.02 using the WillItRebin program(72). The implicit solvent SAXS calculation program Pepsi-SAXS version 3.0 was used to calculate SAXS profiles from the conformational ensemble (i.e., the atomic coordinates of the 10,000 frames) of both peptides (73). The approach, based on the methods described by Larsen et al. and Pesce and Lindorff-Larsen. (34, 74), recommends the parameter values to be set to 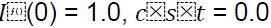, *r*0 = 1.65 Å, and 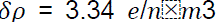. Then, the iterative Bayesian maximum entropy method (34) was used to iteratively rescale and shift the calculated SAXS profiles, while reweighting the conformational ensemble to fit a global value of 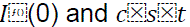.

### NMR

NMR spectra of P3 peptide were recorded on a Bruker Ascend 800 MHz NMR spectrometer (Bruker BioSpin AG, Fälladen, Switzerland) equipped with an Avance NEO console and a 5 mm with cryogenic CP-TCI probe, which is located at the NV-NMR Center at NTNU and is part of the Norwegian NMR Platform (NNP). All NMR recordings were performed on 5 mM peptide in 90:10 H_2_O:D_2_O at 25 °C. For the chemical shift assignment, the following spectra were recorded: 1D proton, 2D in-phase COSY (IP-COSY) (75), 2D total correlation spectroscopy (TOCSY) with 80 ms mixing time, 2D nuclear Overhauser effect correlation spectroscopy (NOESY) with 100 ms mixing time, 2D ^13^C heteronuclear single quantum coherence (HSQC) with multiplicity editing, and 2D ^15^N HSQC. The spectra were recorded, processed and analyzed using TopSpin 4.1 software (Bruker BioSpin) and CcpNMR (76). The chemical shift assignments have been deposited in the BioMagResBank under the accession number 53327. Analysis of secondary chemical shifts was performed on the CSI 3.0 web server (77)

For paramagnetic relaxation enhancement (PRE) in the presence of copper, signal intensities in 2D ^15^N HSQC were measured after the addition of 6 mM CuSO_4_.

### MIC determination

Fungal strains were from the CIRM-CF (Centre International de Ressources Microbiennes-Champignons Filamenteux, INRAE, Marseille, France) collection. Fungi were grown on Potato Dextrose (PD) agar, at 25°C. Fungi suspensions were prepared by scraping spores with sterile water supplemented with NaCl 0.85% and 100 µl L^-1^ Tween 80 and were diluted to 2.5 10^4^ spores mL^-1^ in appropriate broth (RPMI with MOPS or PD for ascomycetes and basidiomycetes, respectively) after counting by microscopy with a calibrated cell. Bacterial suspensions were diluted in MH medium to a final concentration of 10^5^ CFU mL^-1^. *Pco*AA14A_dCTR, P1 or P2 were added to microorganism suspension in polypropylene 96-well microplates at a concentration ranging from 100 to 0.1 µM by 2-fold serial dilutions. Fungi were left for several days to grow at room temperature. MIC was defined as the lowest concentration of peptide inhibiting visible growth. Bacteria were incubated 48h at 37°C. Sterility and growth controls were included in each assay. MICs were determined in three independent replicates.

### Microscopy

P3 and P4 peptides labelled at their N-terminus with 5-carboxyfluorescein (5-FAM; Sigma-Aldrich, #86826), were purchased from GenScript. All peptides were dissolved in Milli-Q ultrapure water and stored at 20 °C. *H. annosum* was grown on potato dextrose agar (PDA; Difco) for two weeks at 25 °C. Spores were collected by gently washing the surface of the plates with sterile water and filtered through sterile Miracloth (Millipore) to remove mycelial fragments and retain dispersed spores. For assays including both spores and hyphae, the Miracloth filtration step was omitted. Spore concentrations were determined using a Neubauer counting chamber*. A. niger* spores were prepared similarly. Spores were suspended at 2 × 10⁶ spores mL^-1^ in 20 mM HEPES, 150 mM NaCl, pH 6.0, prior to mixing with peptides. For green fluorescence binding assays, spores were incubated with 20 μM 5-FAM–labeled P3 peptide. For propidium iodide staining assays, spores were incubated with 250 μM unlabeled peptide, in order to maintain a similar peptide-to-spore ratio as in the MIC experiments. Treatments were carried out in a final volume of 500 μL, with shaking at 450 rpm for 1 hour at 20 °C. In overnight incubation experiments, where hyphal growth was expected, the incubation buffer was supplemented 1:2 with potato dextrose broth (PDB) to support fungal viability. After incubation of peptide–spore mixtures, samples were centrifuged at 10,000 × g for 3 minutes. The supernatant was discarded, and the pellet was washed with 500 μL of Milli-Q water. 480 μL of the supernatant were carefully removed, and the pellet was gently resuspended in the remaining 20 μL of water, yielding a final concentration of 5 × 10⁷ spores mL^-1^ for microscopy imaging. A 10 μL aliquot was transferred onto Quick-Read™ Precision Cell urinalysis slides onto microscopy slides for imaging. To assess the contribution of electrostatic interactions, an additional experiment was performed in which spores were incubated with 20 μM 5-FAM–labeled P3 peptide under identical conditions but in high-salt buffer (20 mM HEPES, 1 M NaCl, pH 6.0). To assess membrane damage, spores treated with peptide or controls were incubated for 1 hour with 0.3 µM propidium iodide prior to microscopy. Propidium iodide does not penetrate intact spores but enters membrane-compromised spores, where it binds to DNA and emits red fluorescence. Microscopy was performed using a 10x or 40x objective in both brightfield and fluorescence modes. Green fluorescence from 5-FAM–labeled peptides was detected using the FITC filter cube, which typically includes an excitation filter around 470–495 nm and an emission filter 510–550 nm. Red fluorescence from propidium iodide was detected using the TRITC filter cube, with an excitation range of 540–570 nm and emission above 590 nm with a fixed exposure time of 500 μs. Image analysis was performed using ImageQuantTL Colony Counter software (Cytiva) to quantify the total number of spores and the number of red-labeled (damaged) spores. All experiments were conducted in at least two biological replicates.

## Acknowledgments

We would like to thank Cendrine Nicoletti (iSm2 lab) and Annick Doan (BBF lab) for their help with the microscopy. We thank Petra Pernot and Mark Tully (ESRF, proposal numbers: MX2324 and MX2490) and Aurélien Thureau (SOLEIL, proposal number: 20210786) for their help in recording the SEC-SAXS data and the ESRF and SOLEIL for synchrotron beamtime allocation. We are also grateful to Gerlind Sulzenbacher (AFMB Lab) for efficiently managing AFMB BAG. We thank all the AFMB technical and support staff (Denis Patrat, Patricia Clamecy, Béatrice Rolland, Chantal Falaschi and Fabienne Amalfitano) and all the BBF support staff (Sabine Genet, Chantal Parodi-Negri, Naura Thibeau and Christophe Boyer). We thank Maria Maté Perez (AFMB Lab) for her help with ITC experiments and SEC-MALLS and Florence Chaspoul (Institut Méditerranéen de Biodiversité et d’Ecologie (IMBE) for ICP-MS analysis. We also thanks Bastien Bissaro for insightful discussions and comments on the manuscript. The authors are grateful to the EPR-MRS facilities of the Aix-Marseille University EPR center and acknowledge the support of the French research infrastructure INFRANALYTICS (FR2054).

This work was carried out with the financial support of the CNRS, INRAE and the French Infrastructure for Integrated Structural Biology (FRISBI) (ANR-10-INSB-0005). K.C.T. was funded by the French government under the France 2030 investment plan, as part of the Initiative d’Excellence d’Aix-Marseille Université (A*MIDEX) and is part of the Institute of Microbiology, Bioenergies and Biotechnology (IM2B) (AMX-19-IET-006). J-G.B., E.T. and K.C. received funding from Campus France in the frame of the PHC Aristote. J-G.B. and G.C. were funded by the Novo Nordisk Foundation (grants NNF20OC0059697 and NNF18OC0032242). G.C. acknowledges the support of the Research Council of Norway through the Norwegian NMR Platform infrastructure grant 226244. Part of the work described was performed using services provided by the 3PE platform, a member of IBISBA-FR (https://doi.org/10.15454/08BX-VJ91; www.ibisba.fr), the French node of the European research infrastructure, EU-IBISBA (www.ibisba.eu). The funders had no role in the design of the study, in the collection, analyses, or interpretation of data, in the writing of the manuscript, or in the decision to publish the results. A CC-BY public copyright license has been applied by the authors to the present document and will be applied to all subsequent versions up to the Author Accepted Manuscript arising from this submission, in accordance with the grant open access conditions.

## Author Contributions

KCT, KC, CR, AL, MH, SG, ML, BG, GC designed and carried out the experiments. KCT, KC, CR, AL, ET, ML, BG, GC, SL, JGB interpreted the data. KT, SL, JGB conceptualized the study. SL, JGB supervised the work and coordinated the study. KT wrote the first draft, and all co-authors contributed with inputs. All authors reviewed and approved the final version of the manuscript.

## Classification

Biological Sciences, Biochemistry

## Supplementary Information

**Figure S1.**
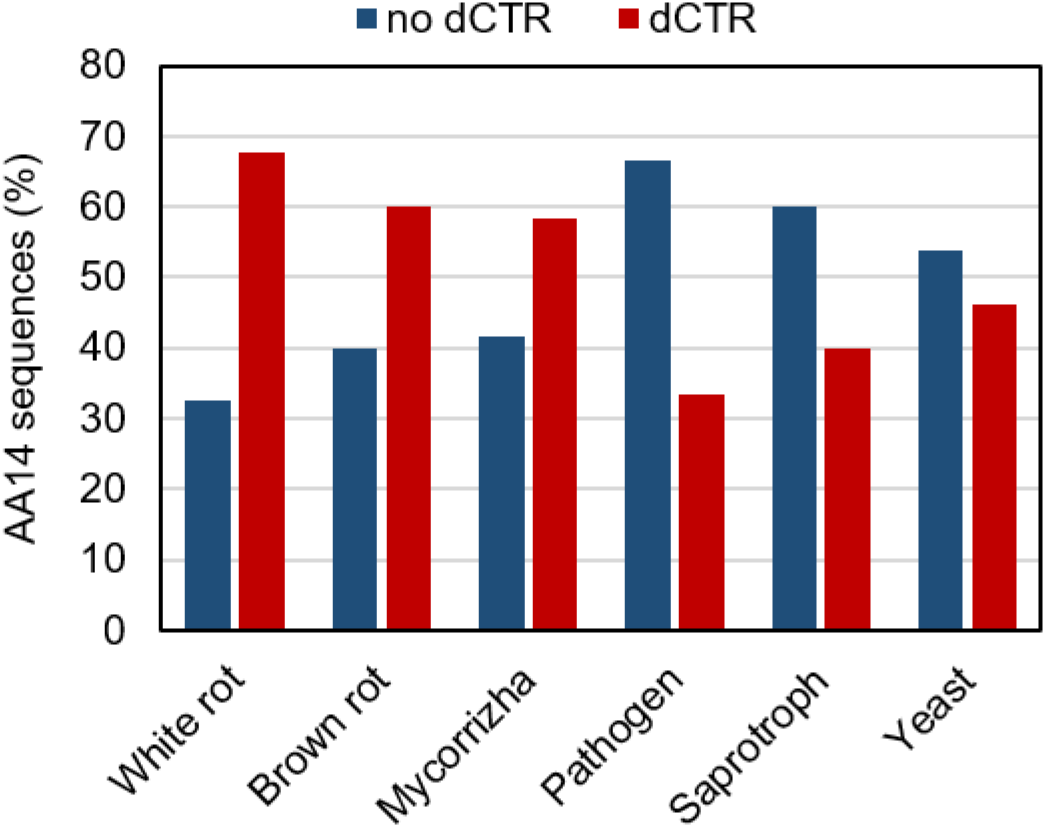
Occurrence of dCTRs in AA14s from organisms with different lifestyles and growth modes.

**Figure S2.**
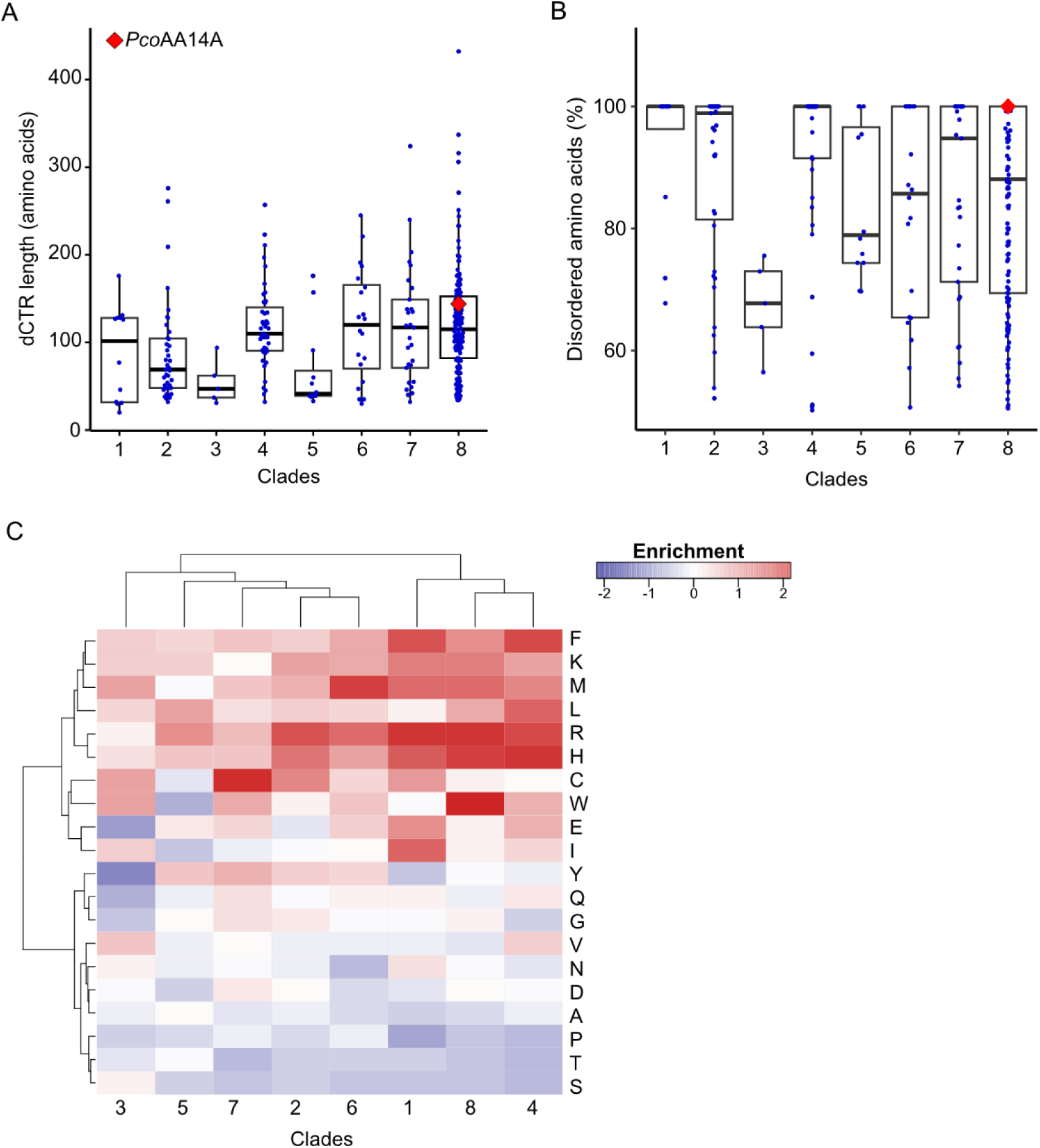
Features of dCTRs from phylogenetically different clades in the AA14 family. (**A**) Boxplots showing the distribution of the length of dCTRs across the different clades in the AA14 LPMO family. (**B**) Boxplots showing the fraction of disordered residues within dCTRs across the clades in the AA14 family. The central box shows the middle portion (i.e., 25–75%) of the dataset: the bottom line of the box defines the first quartile (25%), the middle line shows the median, and the top line of the box shows the third quartile (75%). The extremities of vertical whiskers represent the largest and smallest values. Data points above the upper whiskers represent outliers. The red diamond represents *Pco*AA14A_dCTR. (**C**) Heatmap showing the enrichment or depletion of amino acids found in the last 30 amino acids within dCTRs across different clades.

**Figure S3.**
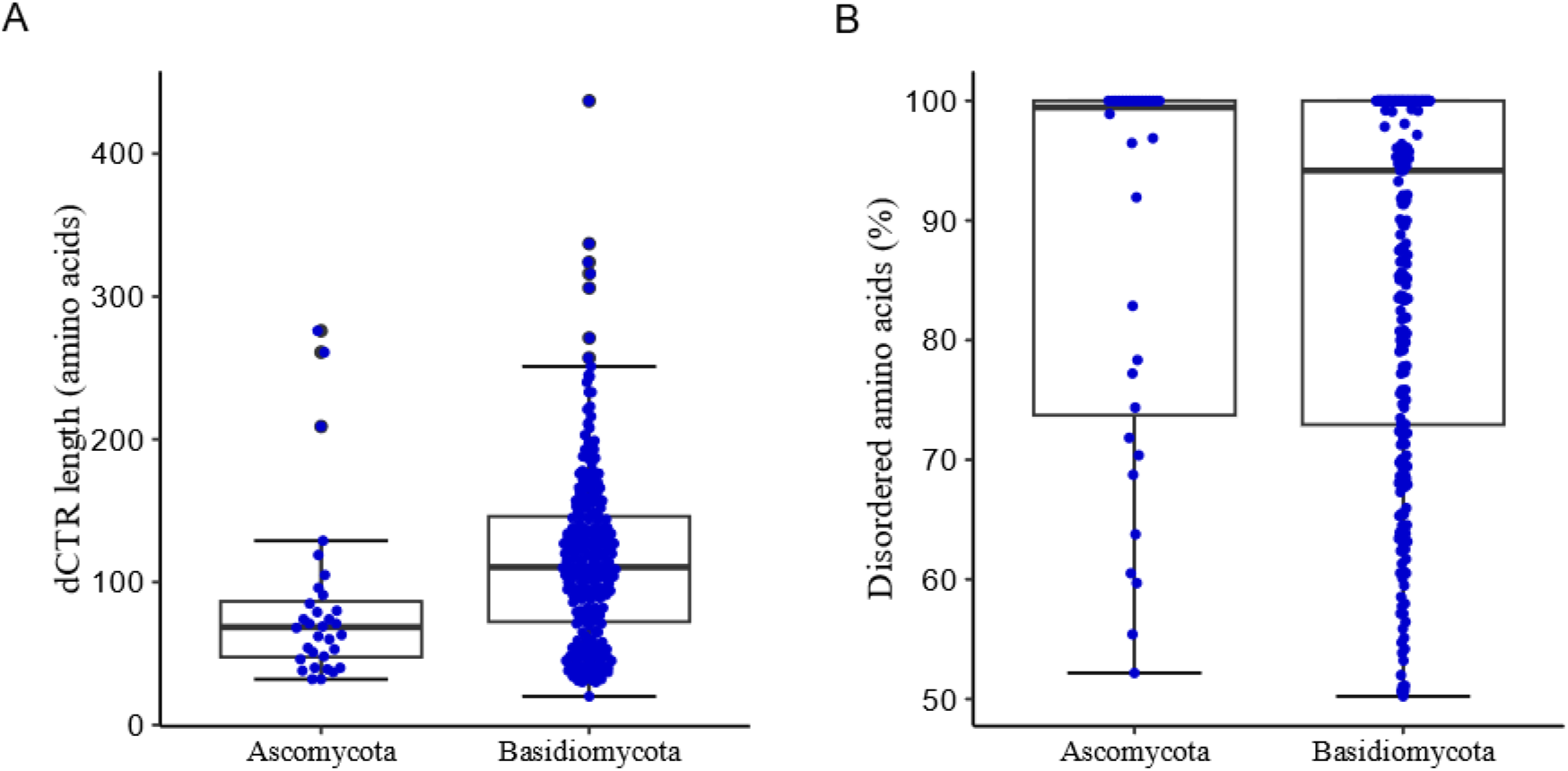
Length (A) and disorder content (B) of AA14-dCTRs from Ascomycetes or Basidiomycetes. The central box shows the middle portion (i.e., 25–75%) of the dataset: the bottom line of the box defines the first quartile (25%), the middle line shows the median, and the top line of the box shows the third quartile (75%). Data points above the upper whiskers represent outliers.

**Figure S4.**
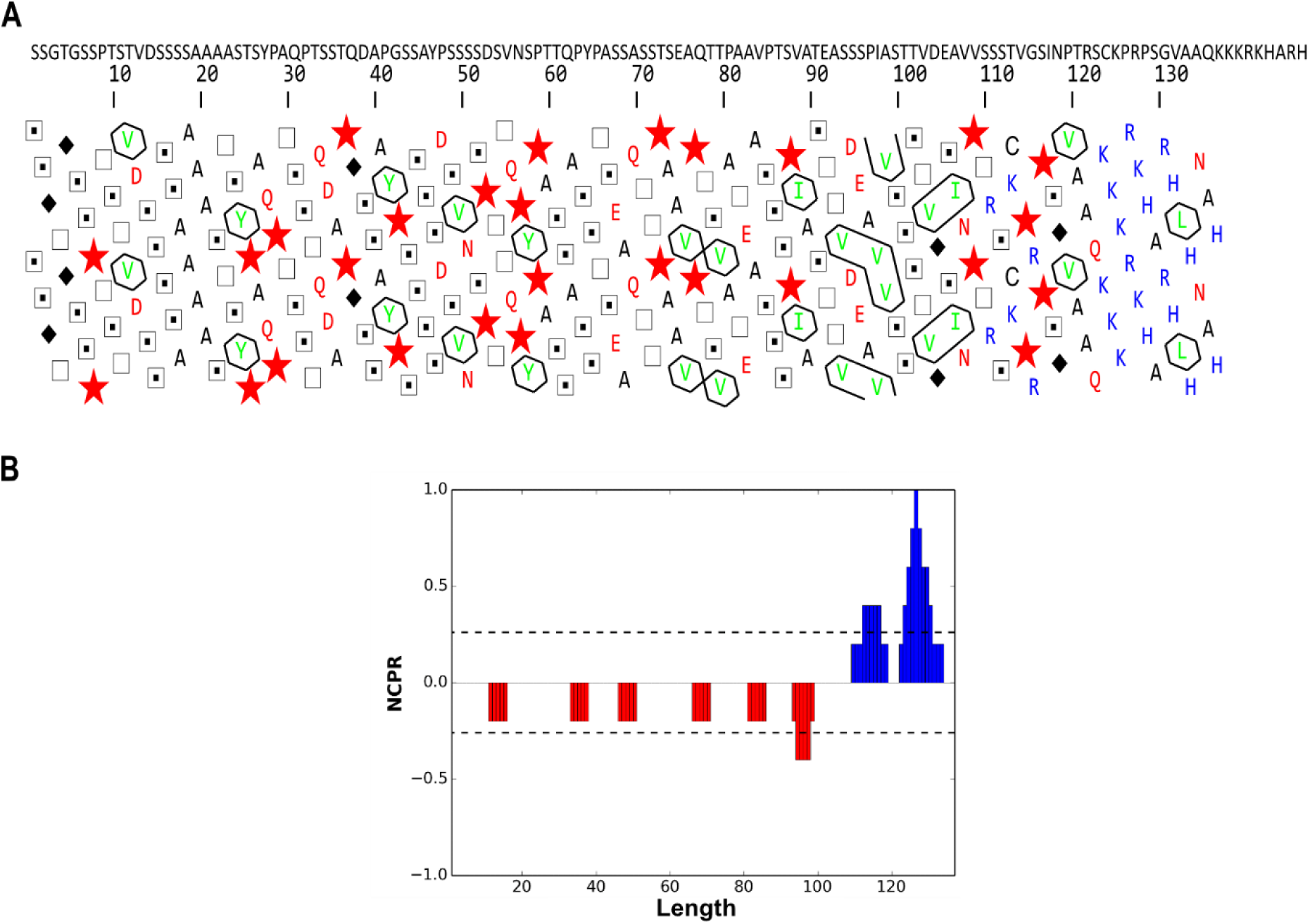
(**A**) Hydrophobic cluster analysis (HCA) plot of *Pco*AA14A_dCTR. Red stars: proline residues, black diamonds: glycine residues, squares: threonine residues, squares with a dot: serine residues. (**B**) Linear net charge per residue (NCPR) of *Pco*AA14A_dCTR as calculated by Cider (Holehouse et al., 2017).

**Figure S5.**
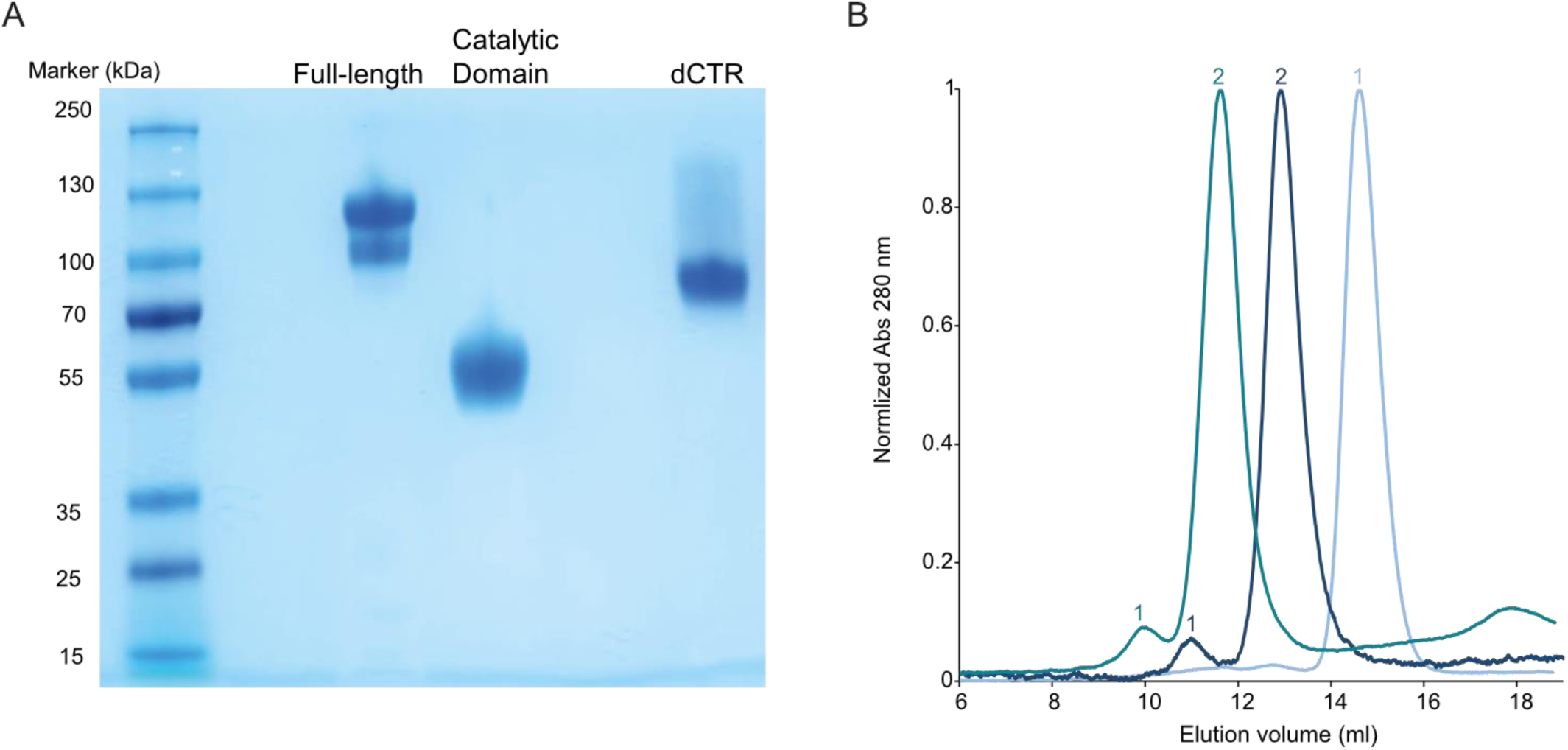
*Pco*AA14A purification. (**A**) SDS-PAGE analysis of purified *Pco*AA14A forms under reducing conditions. (**B**) Size-exclusion chromatography (SEC) of *Pco*AA14A_CD (light blue), *Pco*AA14A_dCTR (blue) and *Pco*AA14A_FL (dark green) (all with a hexahistidine tag) in buffer MES 50mM NaCl 150 mM pH 6.5. The minor (1) and major (2) peaks observed for *Pco*AA14A_dCTR and *Pco*AA14A_FL are labeled.

**Figure S6.**
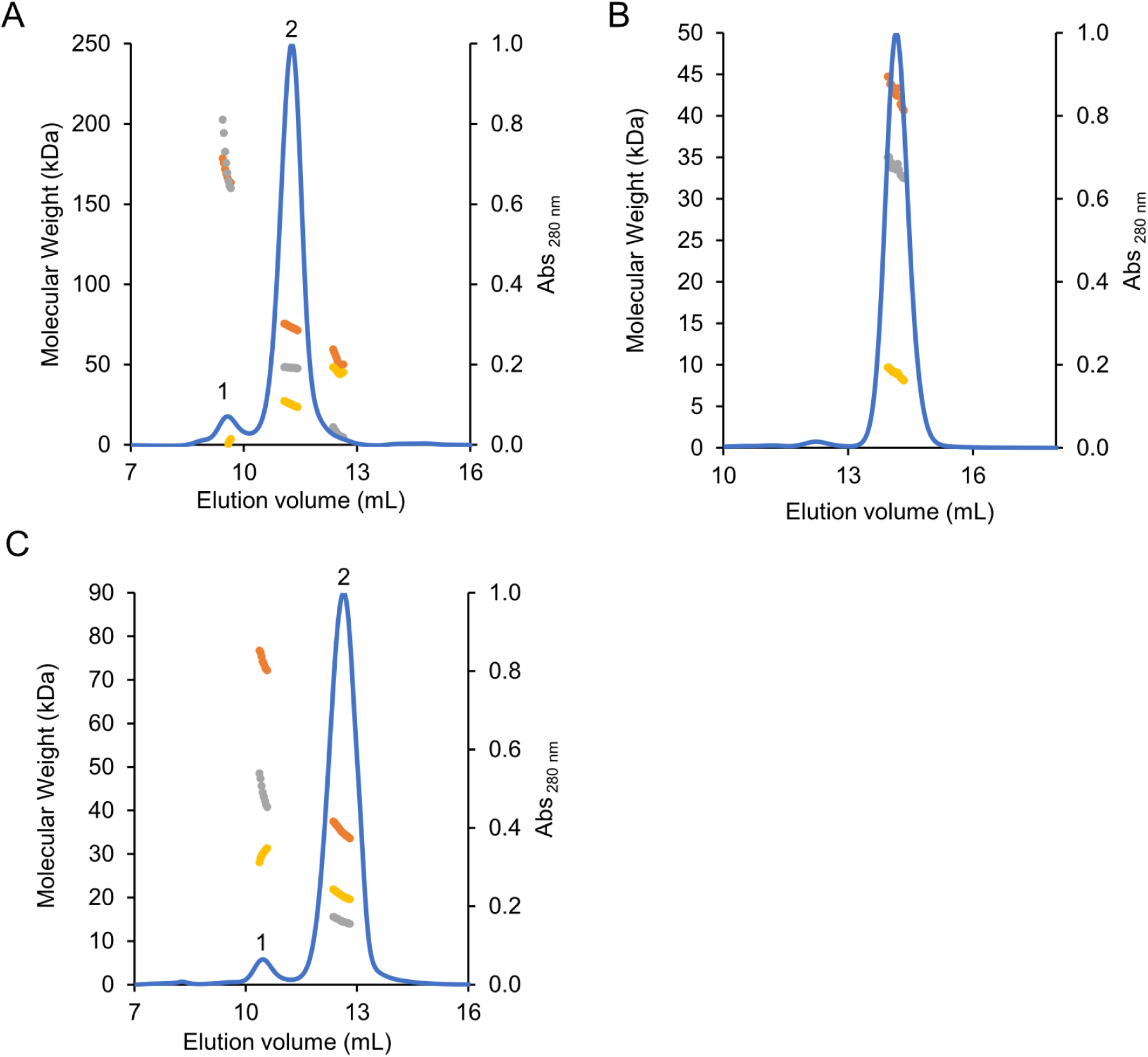
SEC-MALLS elution profiles of (A) *Pco*AA14A_FL, (B) *Pco*AA14A_CD and (C) *Pco*AA14A_dCTR. The orange line corresponds to the total mass, the grey line corresponds to the protein mass and the yellow line corresponds to the glycans mass.

**Table S1.** Molecular mass of *Pco*AA14A_FL, *Pco*AA14A_CD and *Pco*AA14A_dCTR measured by SEC-MALLS. (1) and (2) correspond to the peak #1 and #2 in Figure S6.

|  | *PcoAA14A_FL* | *PcoAA14A_CD* | *PcoAA14A_dCTR* |
| --- | --- | --- | --- |
| Total MM<br>(kDa) | (1) 175<br>(2) 73.5 | 42.4 | (1) 73.8<br>(2) 35.1 |
| Protein MM<br>(kDa) | (1) 170<br>(2) 48 | 33.5 | (1) 29.6<br>(2) 14.6 |
| Sugars MM<br>(kDa) | (1) 5<br>(2) 25.5 | 8.9 | (1) 44.2<br>(2) 20.5 |

**Figure S7.**
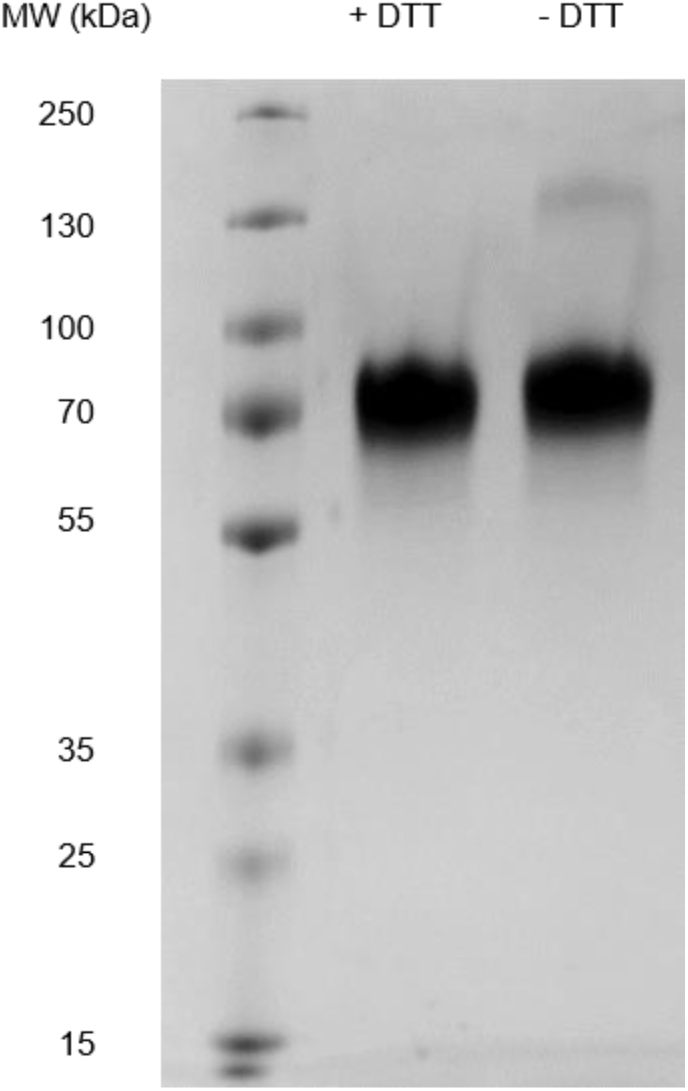
SDS-PAGE analysis of *Pco*AA14A_dCTR in the presence or absence of DTT.

**Figure S8.**
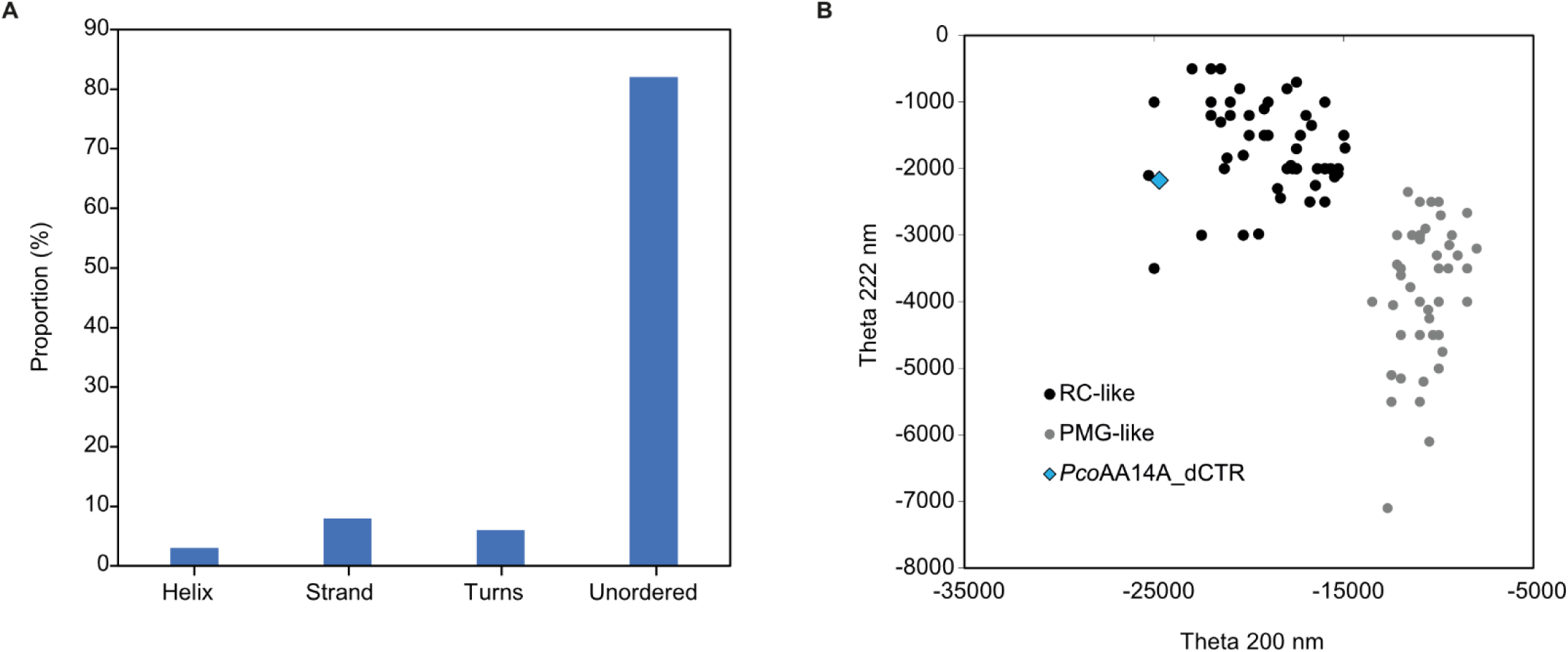
(**A**) Secondary structure content of *Pco*AA14A-dCTR, as derived using Dichroweb (CDSSTR algorithm, set 7). (**B**) Plot of the molar residue ellipticity (MRE) at 222 nm and at 200 nm of a set of well-characterized unfolded, random coil-like (RC-like, black circles) or premolten globule-like (PMG-like, grey circles) proteins (from Uversky et al., 2002). The position in the plot of *Pco*AA14A_dCTR is highlighted in blue.

**Figure S9.**
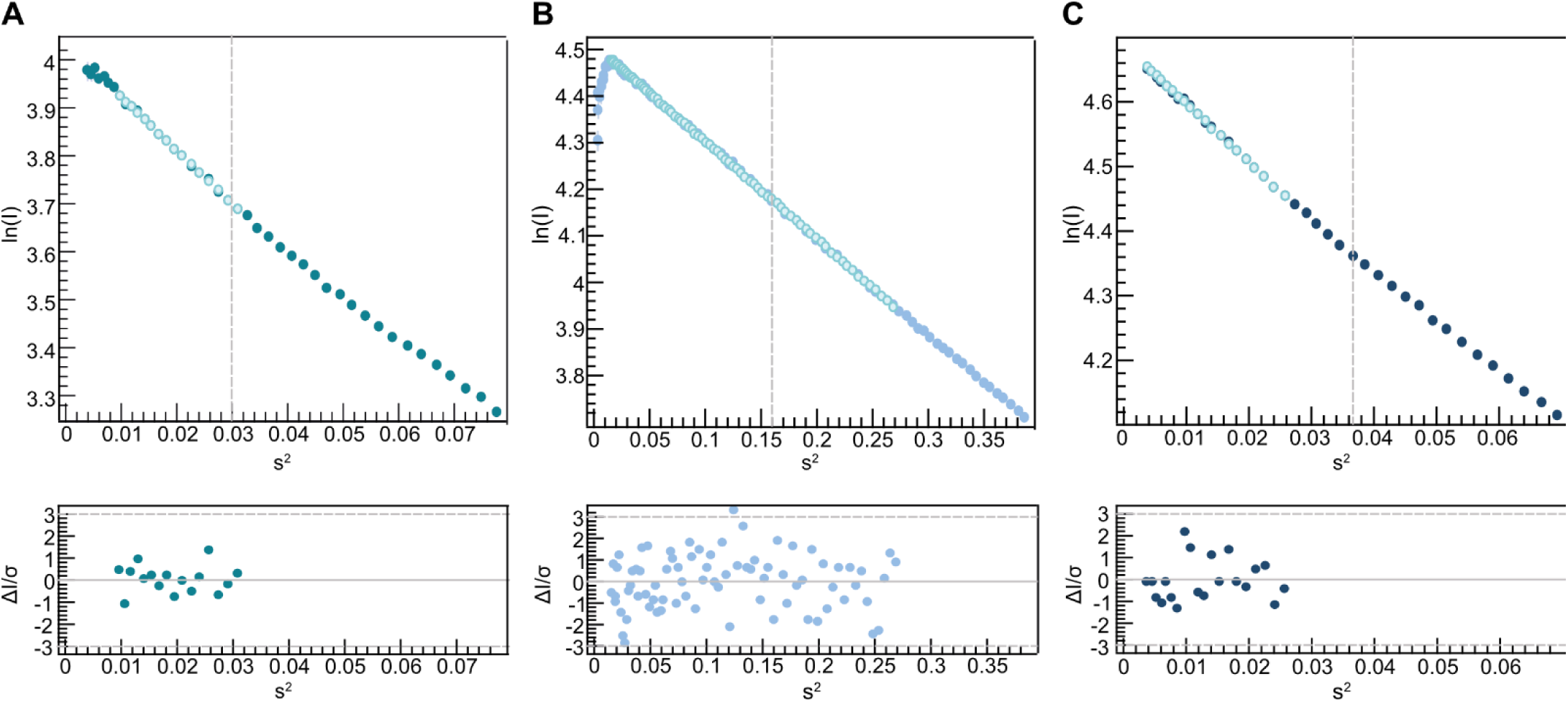
SEC-SAXS analysis. Guinier fitting of (**A**) *Pco*AA14A_FL, (**B**) *Pco*AA14A_CD and (**C**) *Pco*AA14A_dCTR (top panel) scattering curves and plot of residuals (bottom panel).

**Figure S10.**
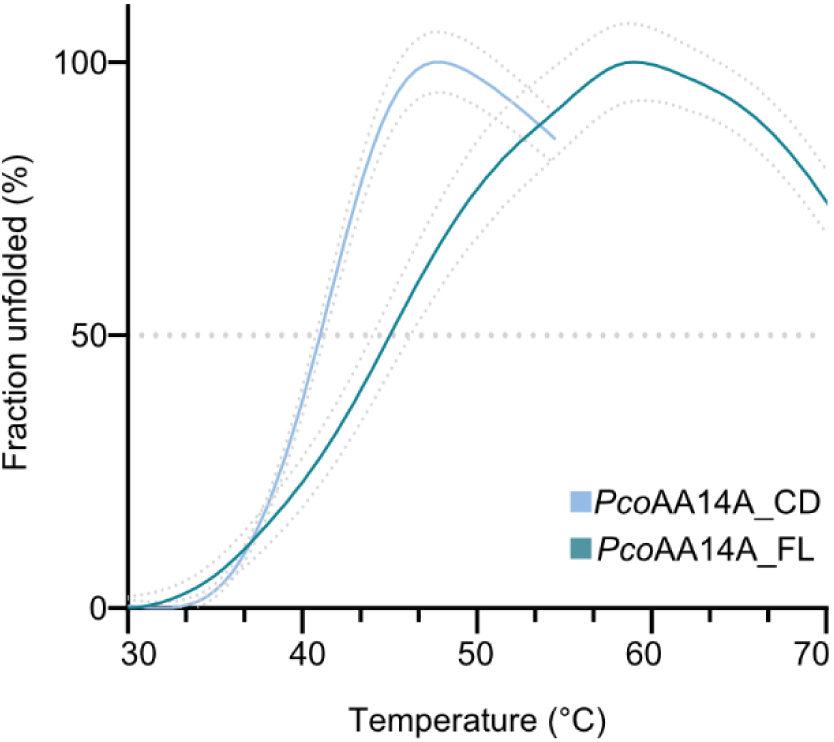
DSF profiles of *Pco*AA14A_FL *and Pco*AA14A_CD. The apparent melting temperatures (T_m_^app^) were determined using a protein thermal shift assay in buffer MES 50 mM NaCl 150 mM pH 6.5. Shown are the average profiles and standard deviations (dots) as obtained from three replicates.

**Figure S11.**
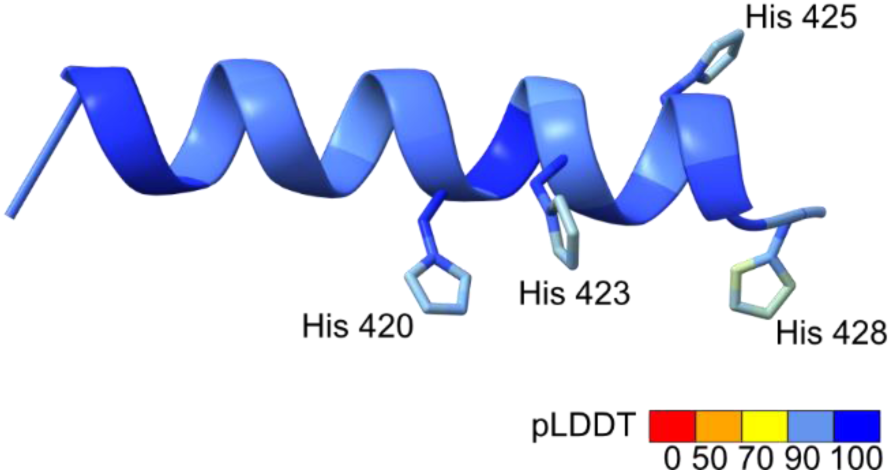
AlphaFold3 model of the last 19 residues of *Pco*AA14A_dCTR showing the His residues potentially involved in copper binding.

**Figure S12.**
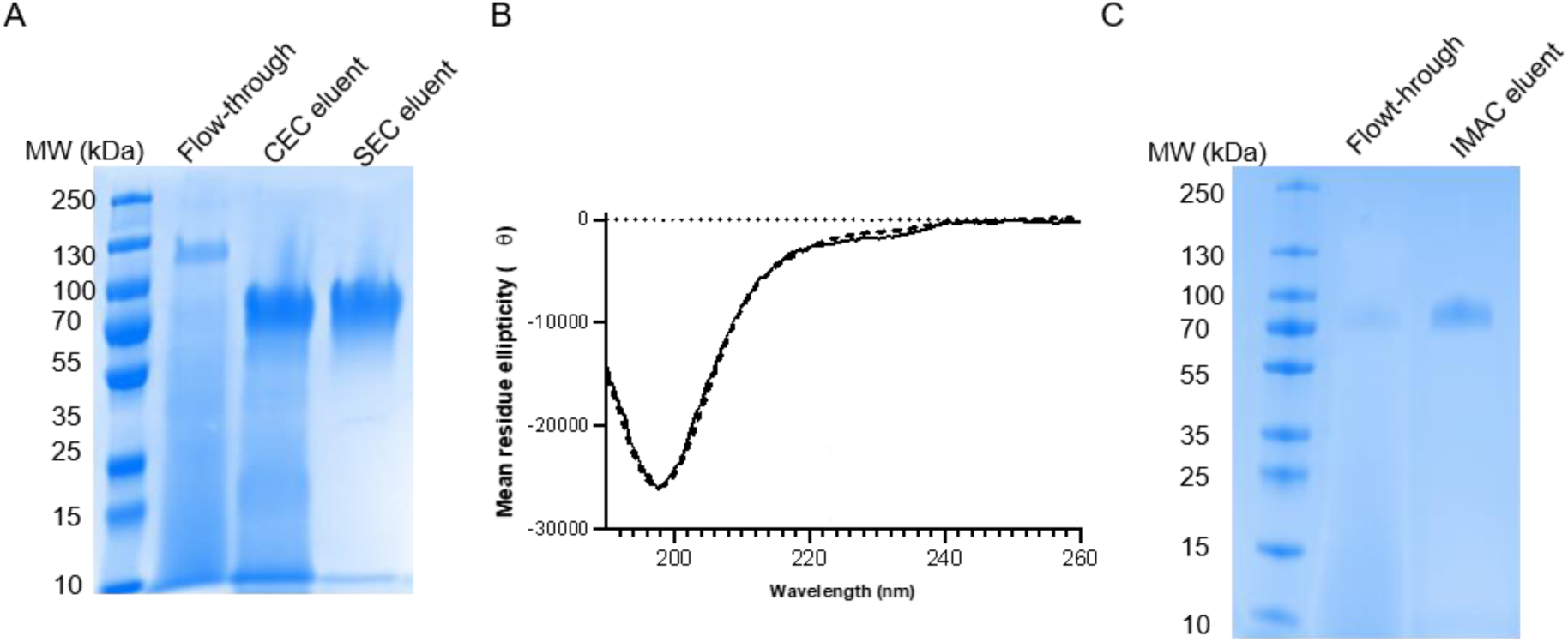
(**A**) SDS-PAGE analysis of the purification of the untagged form of *Pco*AA14A_dCTR. (**B**) Far-UV circular dichroism spectrum of *Pco*AA14A_dCTR with (dashed line) or without (black line) a hexa-histidine tag in 10 mM sodium phosphate pH 6.6. (**C**) SDS-PAGE analysis of the ability of an untagged form of *Pco*AA14A_dCTR to bind to a Histrap column.

**Figure S13.**
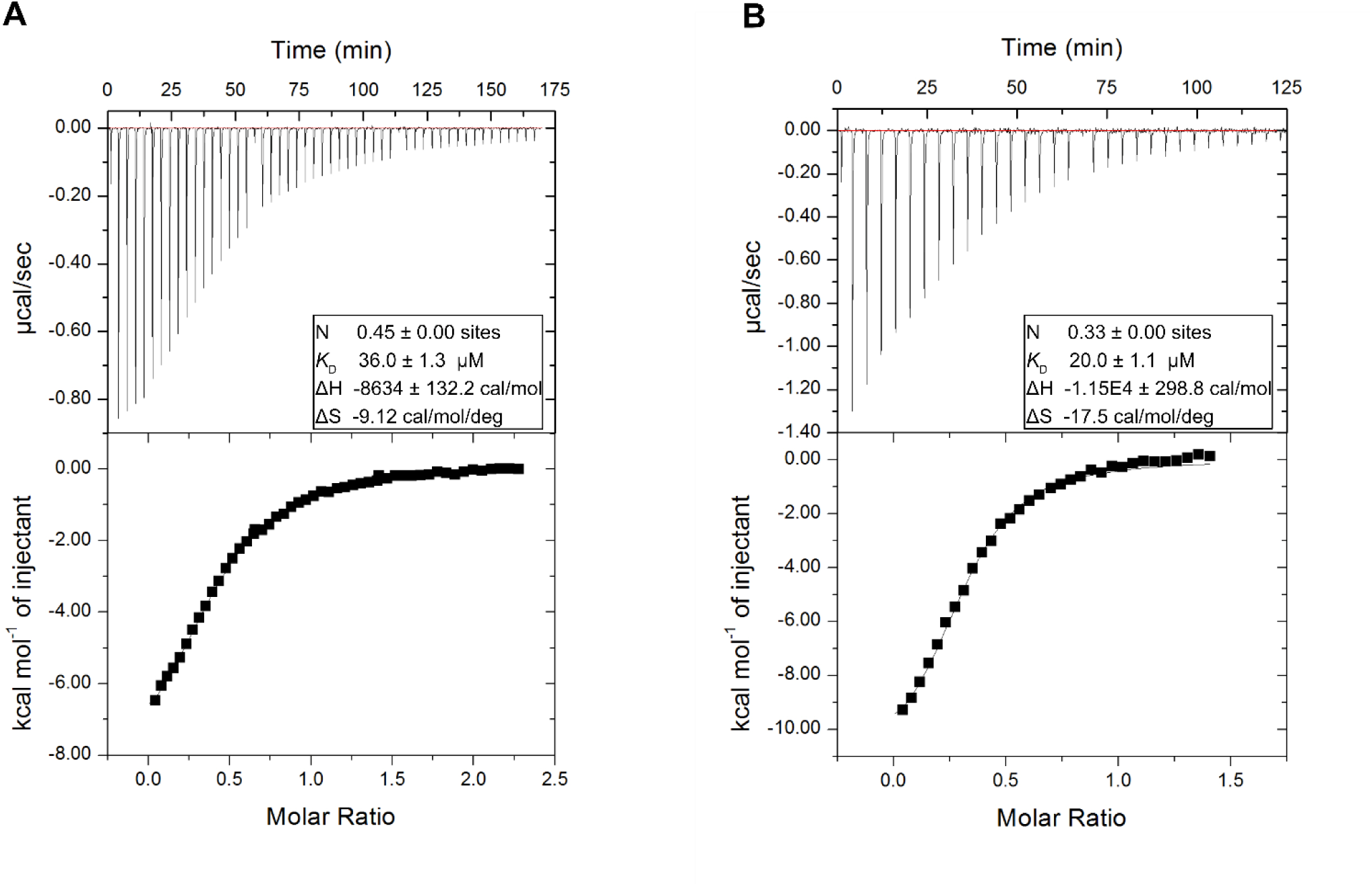
Isothermal titration Calorimetry (ITC) experiment showing the binding of *Pco*AA14A_dCTR (A) and P1 peptide (B) to CuSO_4_. The raw data of the experiment are presented on the top panel, while the binding isotherm for the interaction is shown on the bottom panel. Data of panel A are representative of one out of three independent experiments.

**Table S2.**
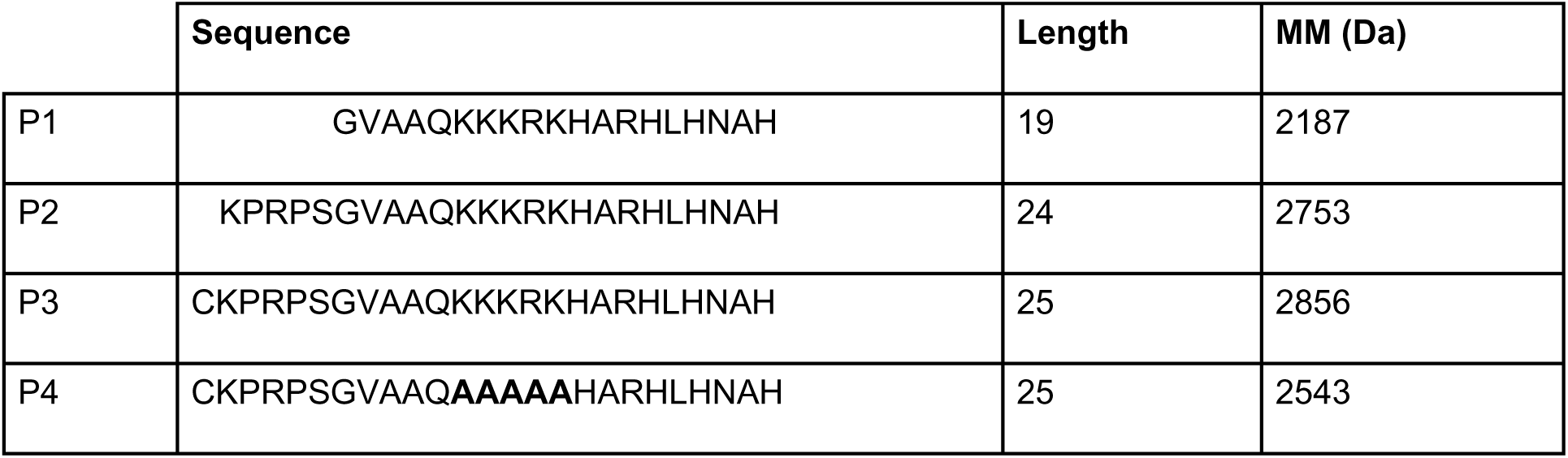
Synthetic peptides used in the study.

**Figure S14.**
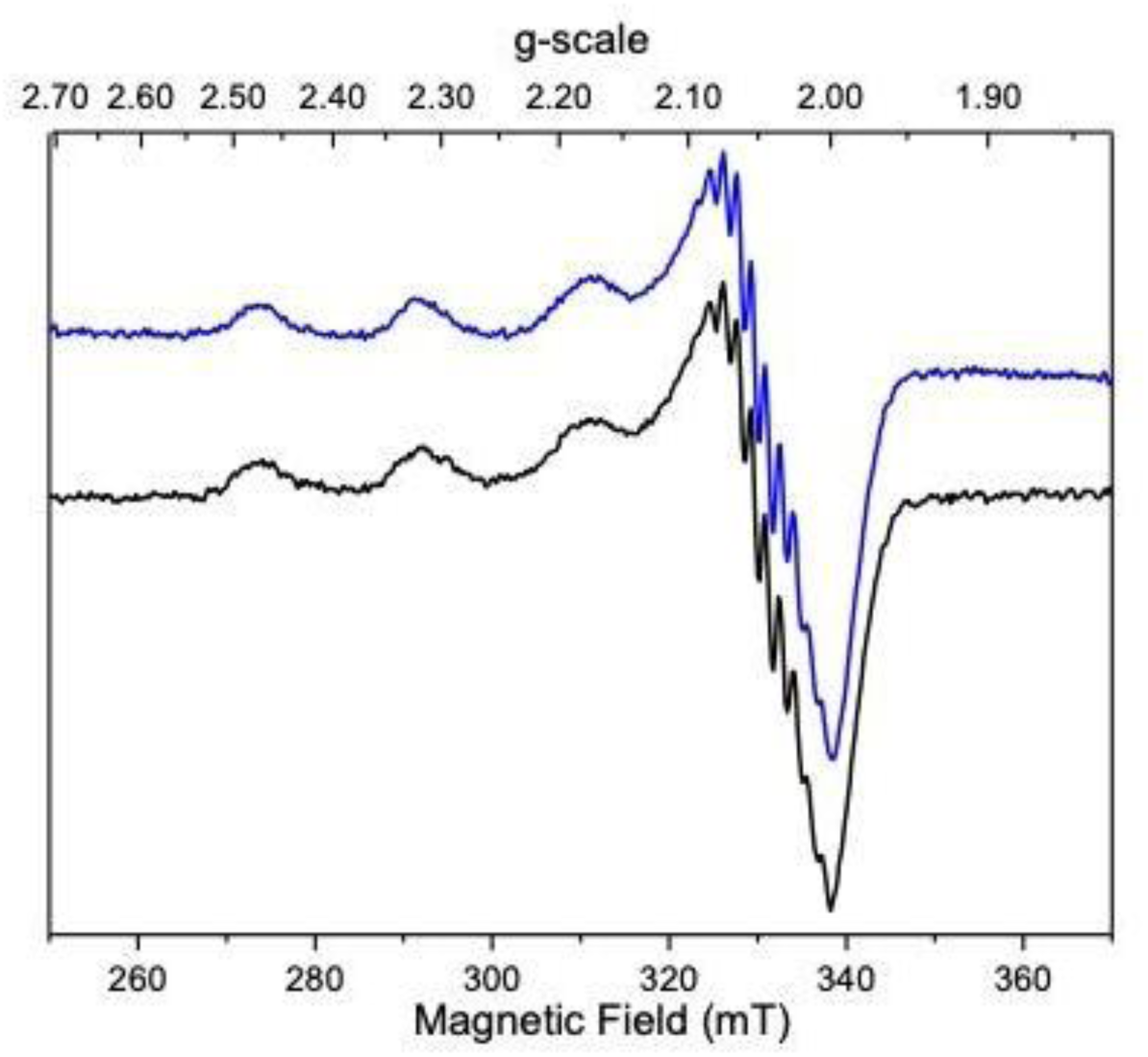
EPR analysis of Cu^2+^ interaction with the *Pco*AA14A dCTR model peptide P1. EPR spectrum of P1 peptide in the presence of 0.5 eq Cu^2+^ at pH 7.2 in HEPES (blue trace) or MOPS (black trace). EPR conditions: temperature, 60 K; microwave power, 4mW at 9.468 GHz; modulation amplitude, 1 mT at 100 kHz.

**Figure S15.**
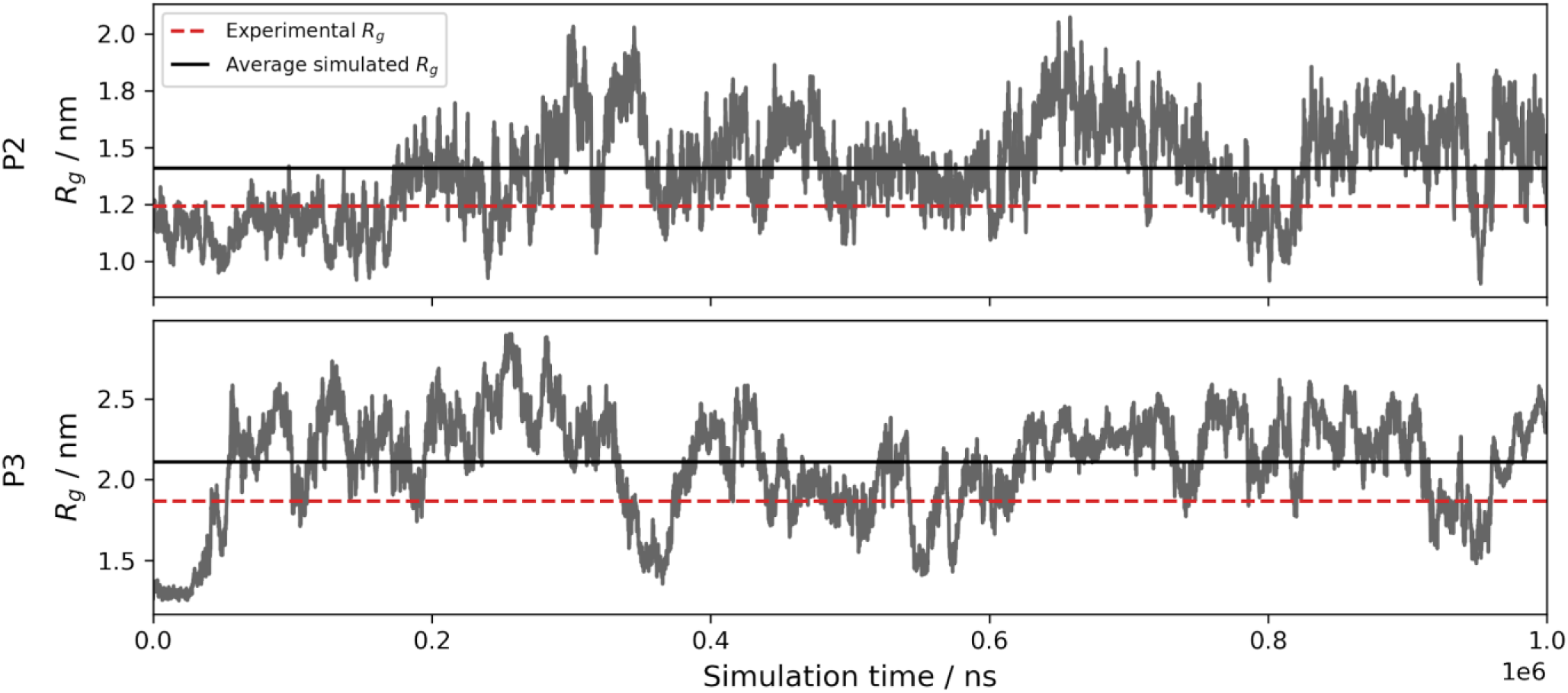
Simulations of P2 and P3 peptides. The graphs show time series of the radius of gyration (R_g_) for simulations of the two peptides. The average simulated R_g_ values (P2=14 Å; P3=21 Å) are shown with solid black lines, and the experimentally determined ones (P2=12.4 ± 0.1 Å; P3=18.7 ± 0.1 Å) are shown by dashed red lines.

**Figure S16.**
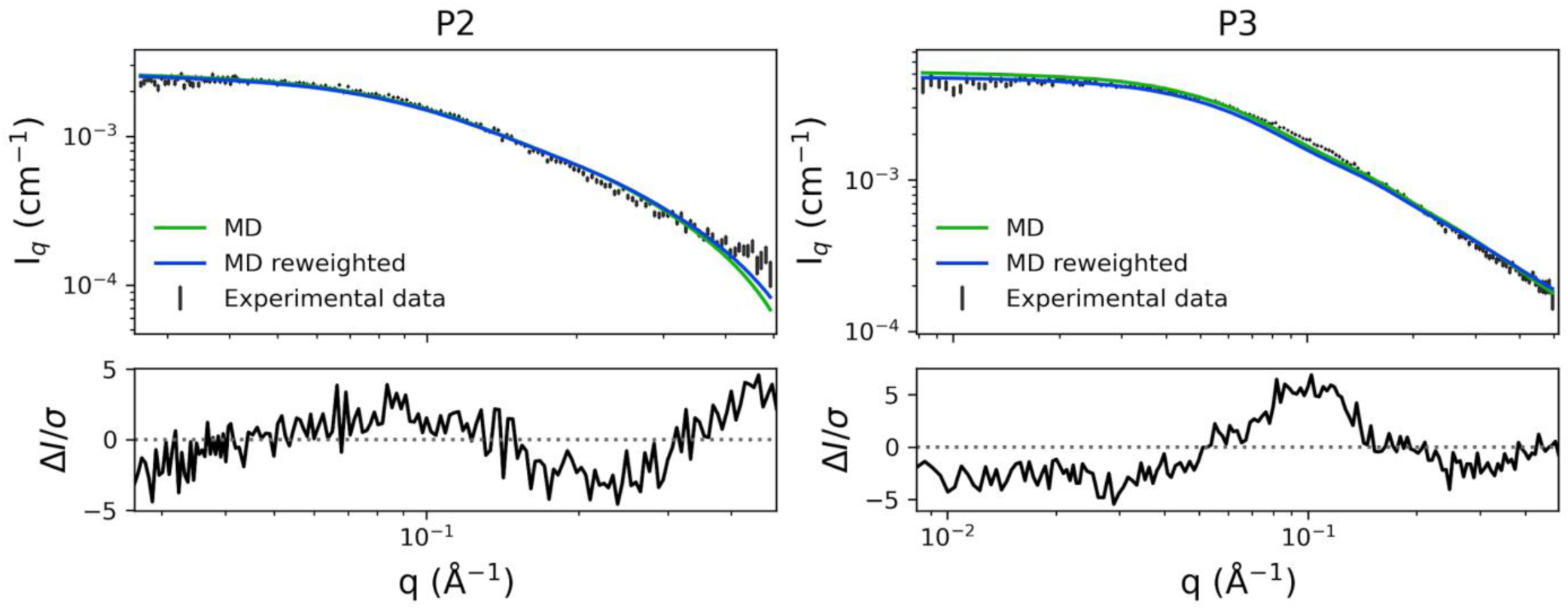
Analysis of the fit to experimental SAXS data of P2 and P3. Experimental SAXS data for P2 and P3 (*black; top panels*) and calculated SAXS data from the ensemble (*green*), and the reweighted ensemble (*blue*). *R*esidual plots are shown on the bottom panels, where ΔI=I_exp_−I_fit_ and σ is the experimental standard deviation. The reason why the experimental and calculated values are different, even when the simulations fit the data well, is because the experimental value includes the size of both the protein and the layer of water around it, while the calculated value only includes the size of the protein.

**Figure S17.**
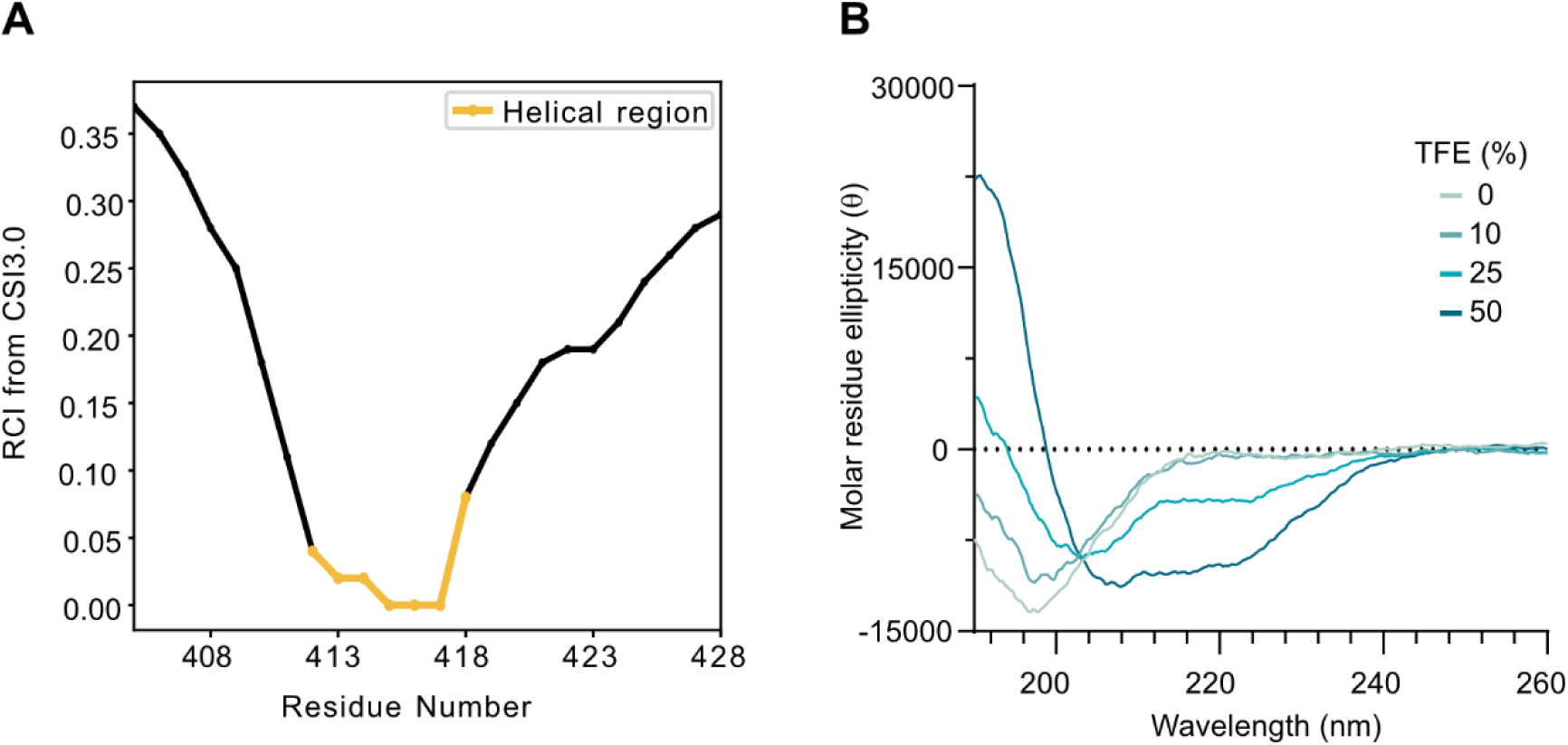
(**A**) Secondary structure propensity of the synthetic peptide P3 determined from the chemical shifts using the Random Coil Index (RCI) from the CSI3.0 web server (Berjanskii and Wishart, 2005; Hafsa et al., 2015). (**B**) Far-UV circular dichroism spectra of the synthetic peptide P2 in the presence of increasing concentration of TFE in 10 mM sodium phosphate pH 6.6.

**Figure S18.**
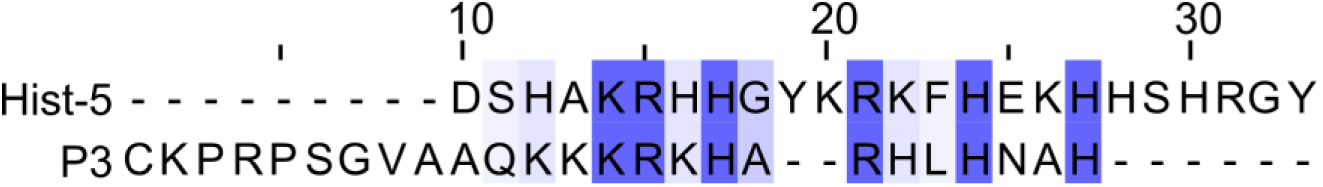
Sequence alignment between Hist-5 and peptide P3.

**Table S3.** Antimicrobial activity of *Pco*AA14A_dCTR. The minimal inhibitory concentration (MIC) obtained with the dCTR, in the presence or absence of copper, and with the P1 and P2 peptides is shown. Each experiment was repeated in three independent replicates. N.D. = not determined.

|  |  |  | MIC (μM) |  |  |  |
| --- | --- | --- | --- | --- | --- | --- |
|  |  |  | dCTR | dCTR + copper | P1 | P2 |
| **Bacteria** | Gram - | *Escherichia coli* | > 100 | > 100 | > 100 | > 100 |
|  |  | *Vibrio alginolyticus* | > 100 | > 100 | >100 | 12.5 |
|  | Gram + | *Bacillus subtilis* | > 100 | > 100 | > 100 | > 100 |
|  |  | Vancomycin-resistant *Enterococcus faecalis* | > 100 | > 100 | > 100 | > 100 |
| **Fungi** | Ascomycetes | *Aspergillus niger* | > 100 | > 100 | > 100 | > 100 |
|  |  | *Penicillium verrucosum* | > 100 | > 100 | > 100 | > 100 |
|  | Basidiomycetes | *Heterobasidion annosum* | 6.25 | 12.5 | 12.5 | 6.25 |
|  |  | *Coniophora puteana* | N.D. | N.D. | 3.12 | 3.12 |

**Figure S19.**
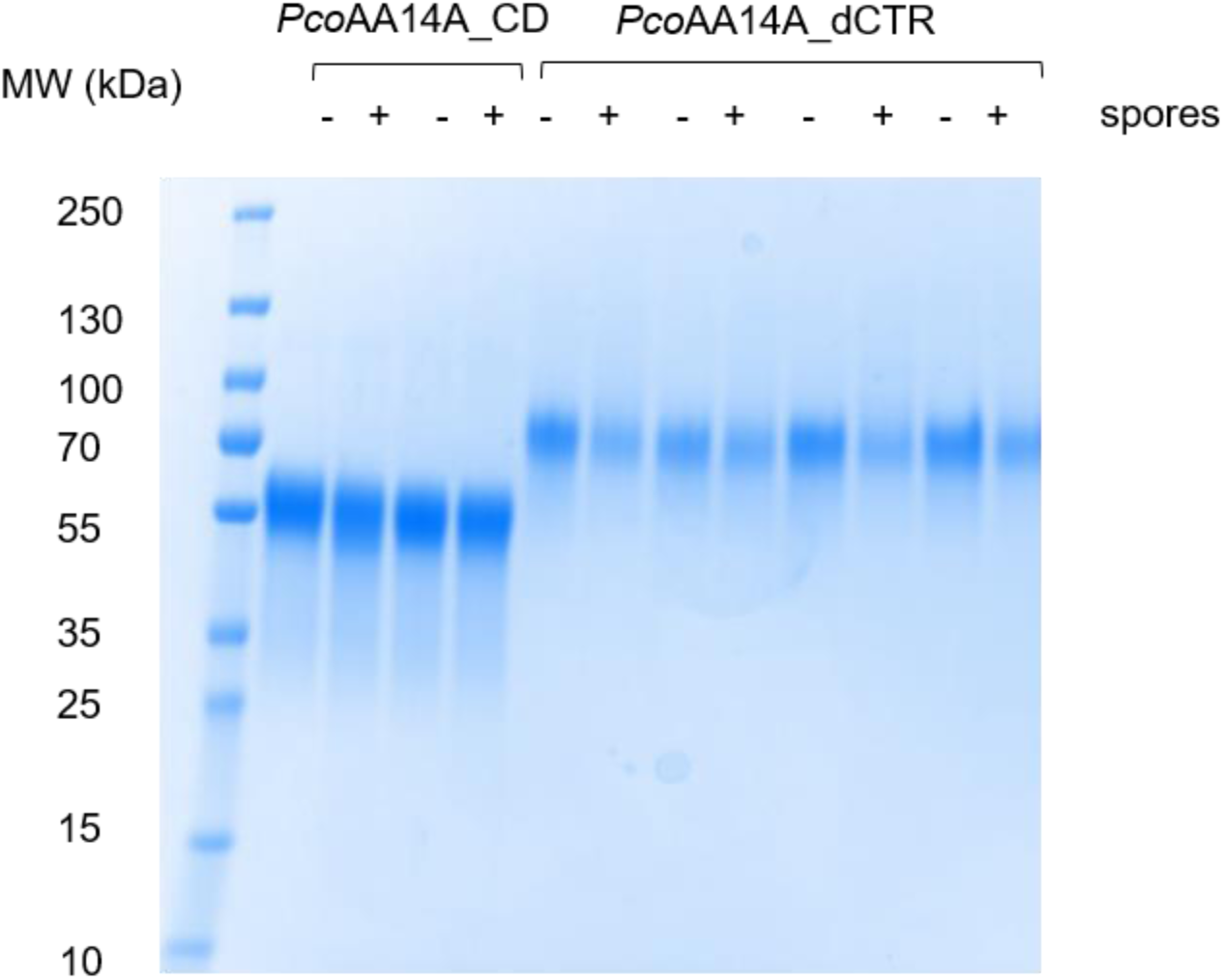
SDS-PAGE analysis of samples of *Pco*AA14A_CD and *Pco*AA14A_dCTR incubated or not with *H. annosum* spores. *Pco*AA14A_CD or *Pco*AA14A_dCTR (30 µM) were incubated with 2 x 10^6^ spores/ml overnight at 25°C. Spores were removed by filtering the samples through a 0.22 µm filter, and the filtrate was loaded onto an SDS-PAGE gel. Multiple lanes correspond to two independent experiments for *Pco*AA14A_CD and four independent experiments for *Pco*AA14A_dCTR.

**Figure S20.**
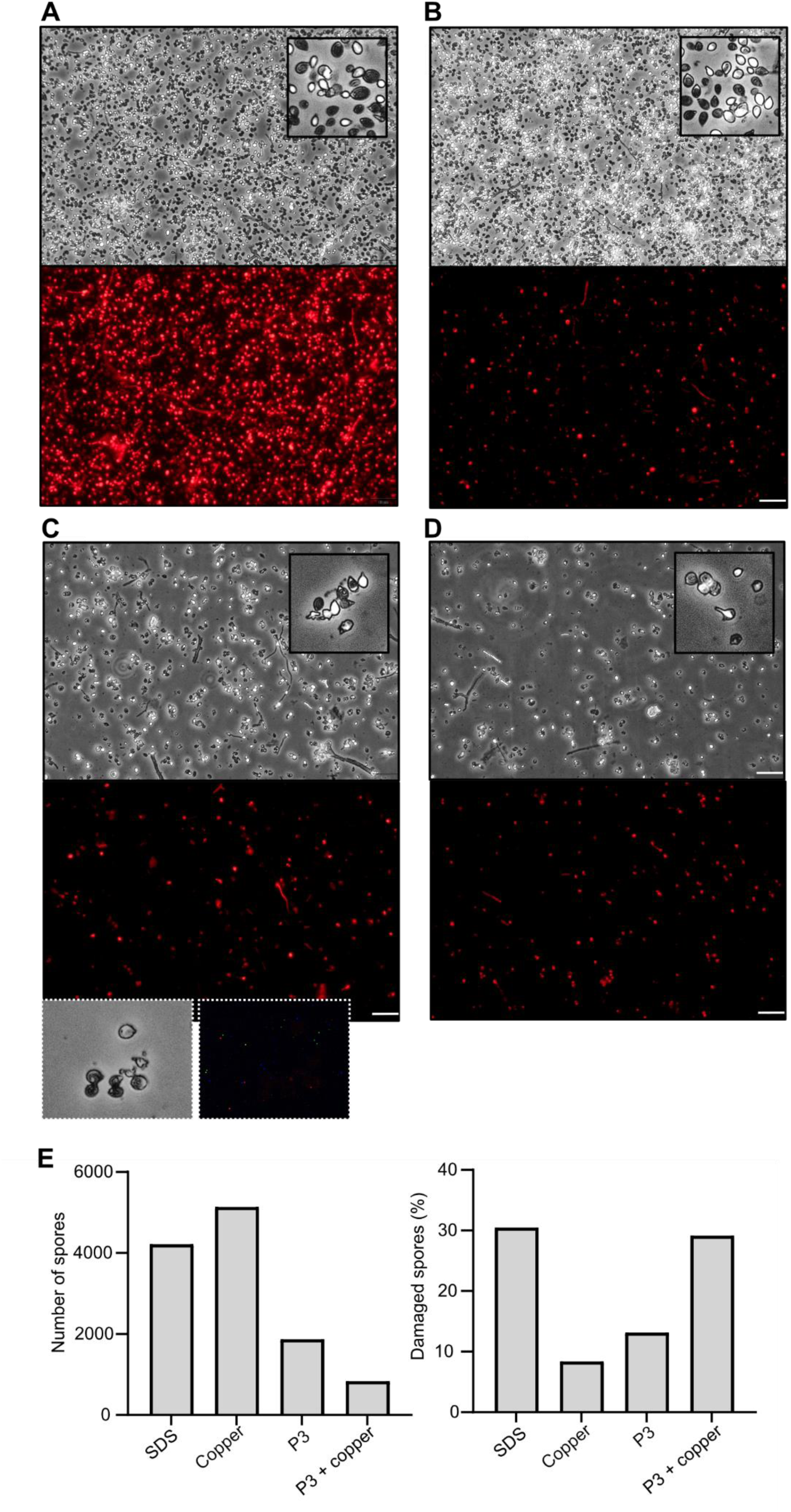
Fluorescence microscopy analysis of propidium iodide labeling of *H. annosum spores* incubated with P3 or P4 peptide. (**A**) Spores were treated with 0.2% SDS, serving as a positive control for chemically induced cell damage. The upper panel shows brightfield microscopy, displaying the total spore population. The lower panel shows fluorescence microscopy using propidium iodide to detect red-labeled, membrane-compromised (damaged) spores. (**B**) Brightfield (top) and fluorescence (bottom) images of spores treated with 250 μM CuSO₄ (negative control). (**C**) Brightfield (top) and fluorescence (bottom) images of spores treated with 250 μM P3 peptide. Inset: spores treated with a P4 mutated peptide lacking charged amino acids. No detectable red fluorescence was observed. (**D**) Brightfield (top) and fluorescence (bottom) images of spores treated with both 250 μM P3 peptide and 250 μM CuSO₄. (**E**) Total number of spores (left) and percentage of damaged spores (right) in tested conditions. The overall number of spores was reduced by 63.7% and 83.9% in the presence of P3 and P3 + CuSO₄, respectively, suggesting that the treatments may lead to spore loss due to aggregation, damage, or other factors affecting visibility or recovery. All treatments were performed at 20 °C for one hour in the presence of propidium iodide, followed by microscopy analysis. Experiments were conducted in at least two independent replicates. Scale bars: 30 μm.

